# Characterization of Peripheral Blood Mononuclear Cells Gene Expression Profiles of Pediatric *Staphylococcus aureus* Persistent and Non-Carriers Using a Targeted Assay

**DOI:** 10.1101/544494

**Authors:** Elisabeth Israelsson, Damien Chaussabel, Rebecca S.B. Fischer, Heather C. Moore, D. Ashley Robinson, Heather T. Essigmann, Eric L. Brown

**Affiliations:** Department of Systems Immunology, Benaroya Research Institute at Virginia Mason, Seattle, WA, USA; Division of Epidemiology, Human Genetics, and Environmental Sciences, University of Texas Health Science Center, Houston, TX, USA; University of Texas McGovern Medical School, Northwest Assistance Ministries, Houston, TX, USA; Department of Microbiology and Immunology, University of Mississippi Medical Center, Jackson, MS, USA

**Author notes:** Current affiliation: Target & Translational Science Department, Respiratory, Inflammation & Autoimmunity, Innovative Medicines and Early Development Biotech Unit, AstraZeneca, Gothenburg, Sweden (Elisabeth Israelsson). Current affiliation: Systems Biology Department, Sidra Medical and Research Center, Doha, Qatar (Damien Chaussabel). Current affiliation: Texas A&M Health Science Center School of Public Health, Department of Epidemiology and Biostatistics, College Station, Texas, USA. Current affiliation: Baylor College of Medicine, Complex Care Clinic, Texas Children’s Hospital, Houston, Tx, USA. Address correspondence to Eric L. Brown.

**Keywords:** *Staphylococcus aureus*, targeted assay, gene expression, RNAseq, carriage, genetics, innate immunity

## Abstract

Defects in innate immunity affect many different physiologic systems and several studies of patients with primary immunodeficiency disorders demonstrated the importance of innate immune system components in disease prevention or colonization of bacterial pathogens. To assess the role of the innate immune system on nasal colonization with *Staphylococcus aureus*, innate immune responses in pediatric *S. aureus* nasal persistent carriers (n=15) and non-carriers (n=15) were profiled by analyzing co-clustered gene sets (modules) identified through large-scale transcriptome data analysis as the basis for the development of a targeted assay. We stimulated previously frozen peripheral blood mononuclear cells (PBMCs) from these subjects with i) a panel of TLR ligands, ii) live *S. aureus* (either a mixture of strains or stimulation with respective carriage isolates), or iii) heat-killed *S. aureus*. We found no difference in responses between carriers and non-carriers when PBMCs were stimulated with a panel of TLR ligands. However, PBMCs stimulated with live *S. aureus* elicited a significantly different response that also differed from the response elicited following stimulation with dead *S. aureus*. Furthermore, we observed a distinct stimulation profile for PBMCs isolated from persistent carriers stimulated with their respective live or dead carriage strains compared to responses observed for PBMCs isolated from non-carriers that were similar regardless of whether or not the bacteria were alive or not. These data suggested that innate pathway signaling is different between persistent and non-carriers of *S. aureus*.

## Introduction

The spectrum of diseases resulting from *S. aureus* infections includes serious skin infections, endocarditis, arthritis, osteomyelitis, and sepsis as a consequence of its ability to colonize a variety of different tissues and its ability to circumvent various immune surveillance systems (1–12). Approximately 20-50% of adults and children in the United States are nasal *S. aureus* carriers (persistent or intermittent carriers vs. non-carriers) based on the presence or absence of *S. aureus* in nasal cultures collected over time (13–16). Nasal sampling over a defined time period allows classification of *S. aureus* carrier phenotypes as either persistent carriers (those testing positive ≥75% of the time), intermittent carriers (individuals <75% of the time) and non-carriers (negative for *S. aureus* over the sampling period) (17). Although a genome-wide association study conducted by our group suggests that each carriage phenotype is in part shaped by host genetic profiles (16), characterization of antibody responses and *S. aureus* re-colonization studies suggest that the intermittent and non-carrier phenotypes are more similar to each other compared to persistent carriers (18, 19).

A combination of environmental and genetic factors play roles in defining the respective *S. aureus* carriage phenotypes and candidate gene studies have linked specific genes with persistent *S. aureus* nasal carriage. For example, polymorphisms in immune-related genes (IL-4, mannose binding lectin [complement pathway], C1 inhibitor [complement pathway], complement factor H, C-reactive protein, vitamin D receptor polymorphisms, β-defensin production, and Toll-like receptor 2) have been shown to contribute to persistent *S. aureus* nasal carriage (20–27).

Despite these insights into immune components that promote persistent *S. aureus* carriage, no effective human vaccine that prevents colonization or infections has been developed (28). This failure can be partially explained by our lack of understanding regarding the nature of the immune response(s) required to prevent *S. aureus* infections/colonization. Due to the clear differences in anti-*S. aureus* immune responses observed between persistent and intermittent/non-carriers (18, 19) we examined the role of innate immune responses using high-throughput methods for transcript profiling. Our goal was to provide a better understanding of which innate immune gene modules are associated with these distinct carriage profiles in a pediatric cohort.

The successful assessment of innate immune responsiveness to respective pathogens could result in the development of novel vaccines and new diagnostic tools for patients presenting with recurrent and/or severe infections. We have previously reported on the design of modules consisting of sets of coordinately expressed transcripts (29). The modular framework has been successfully implemented in a wide range of studies, covering infections and autoimmune diseases (30–32). Recently, we reported on the creation of a set of modules designed to further investigate gene profiles associated with innate immune activation signals (33). The resulting 66 modules are divided into 7 clusters based on their response specificities; some are specific for bacterial stimuli while others are specific for stimuli via a TIR (Toll-IL-1 receptor) domain.

Here we make use of the dimension-reduction concept we have developed for microarray analysis and use it for gene selection for a focused qPCR-based assay in order to develop a time- and cost-efficient tool that can be used to assess potential susceptibilities in individual patients or groups. We used this targeted tool kit for the assessment of innate immune response profiles of peripheral blood mononuclear cells (PBMCs) isolated from pediatric *S. aureus* persistent and non-carriers stimulated with either i) a panel of TLR ligands, ii) live *S. aureus*, or iii) heat-killed *S. aureus*. This analysis demonstrates that children who are persistently colonized with *S. aureus* displayed a unique innate immune response profile to *S. aureus* stimulation compared to non-carrier controls. Furthermore, the immune response elicited in response to their specific *S. aureus* carriage strain was distinct from responses elicited by other *S. aureus* strains. No difference in responses between carriers and non-carriers were observed when PBMCs were stimulated with a panel of TLR ligands.

## Results

### Pediatric *S. aureus* carriage study population

Five persistent carriers had prior medical histories of culture-positive *Staphylococcal* infections compared to non-carriers that lacked any infection history (p=0.042) (Table 1). In addition, MRSA (methicillin resistant *S. aureus*) persistent carriers were more likely to have developed an *S. aureus* infection than MSSA (methicillin susceptible *S. aureus*) persistent carriers (75% vs. 9%, respectively, p=0.033). Most persistent carriers harbored MSSA (n=12; 80%) and 3 (20%) were MRSA persistent carriers. *S. aureus* isolates recovered from 12 persistent carriers were available for our stimulation assay (Table 2). PBMCs harvested from these 12 carriers were stimulated with their respective live carriage isolate. In addition, the 12 isolates were then combined to form the ‘Carrier Mix’ for the stimulation of PBMCs (Table 2).

**Table 1.** Characteristics of the Study Population (n=30).

| **Table 1**<br>Characteristics of the Study Population (n=30). |  |  |  |
| --- | --- | --- | --- |
|  | **Non-Carriers**<br>15 (50%) | **Persistent Carriers**<br>15 (50%) | p-value |
| Age (years) |  |  |  |
| <6 | 6 (40%) | 7 (47%) | 1.000 |
| ≥6 | 9 (60%) | 8 (53%) |  |
| Sex |  |  |  |
| Female | 7 (47%) | 5 (33%) | 0.710* |
| Male | 8 (53%) | 10 (67%) |  |
| Race/Ethnicity |  |  |  |
| Caucasian, non-Hispanic | 0 (0%) | 2 (13%) | 0.636 * |
| Latino/Hispanic | 12 (80%) | 9 (60%) |  |
| African American | 2 (13%) | 3 (20%) |  |
| Other | 1 (7%) | 1 (7%) |  |
| History of Culture-Positive Staphylococcal Infection |  |  |  |
| No | 15 (100%) | 10 (67%) | 0.042 †‡ |
| Yes | 0 (0%) | 5<sup>§</sup> (33%) |  |
| Number of Nasal Swabs Completed |  |  |  |
| 3 | 10 (67%) | 13 (87%) | 0.390 |
| 4 | 5 (33%) | 2 (13%) |  |
| *Fisher's exact p-value. |  |  |  |
| †Statistically significant at p<0.05. |  |  |  |
| §Four patients had MRSA infections; 3 of those patients were MRSA persistent carriers. |  |  |  |

**Table 2.**
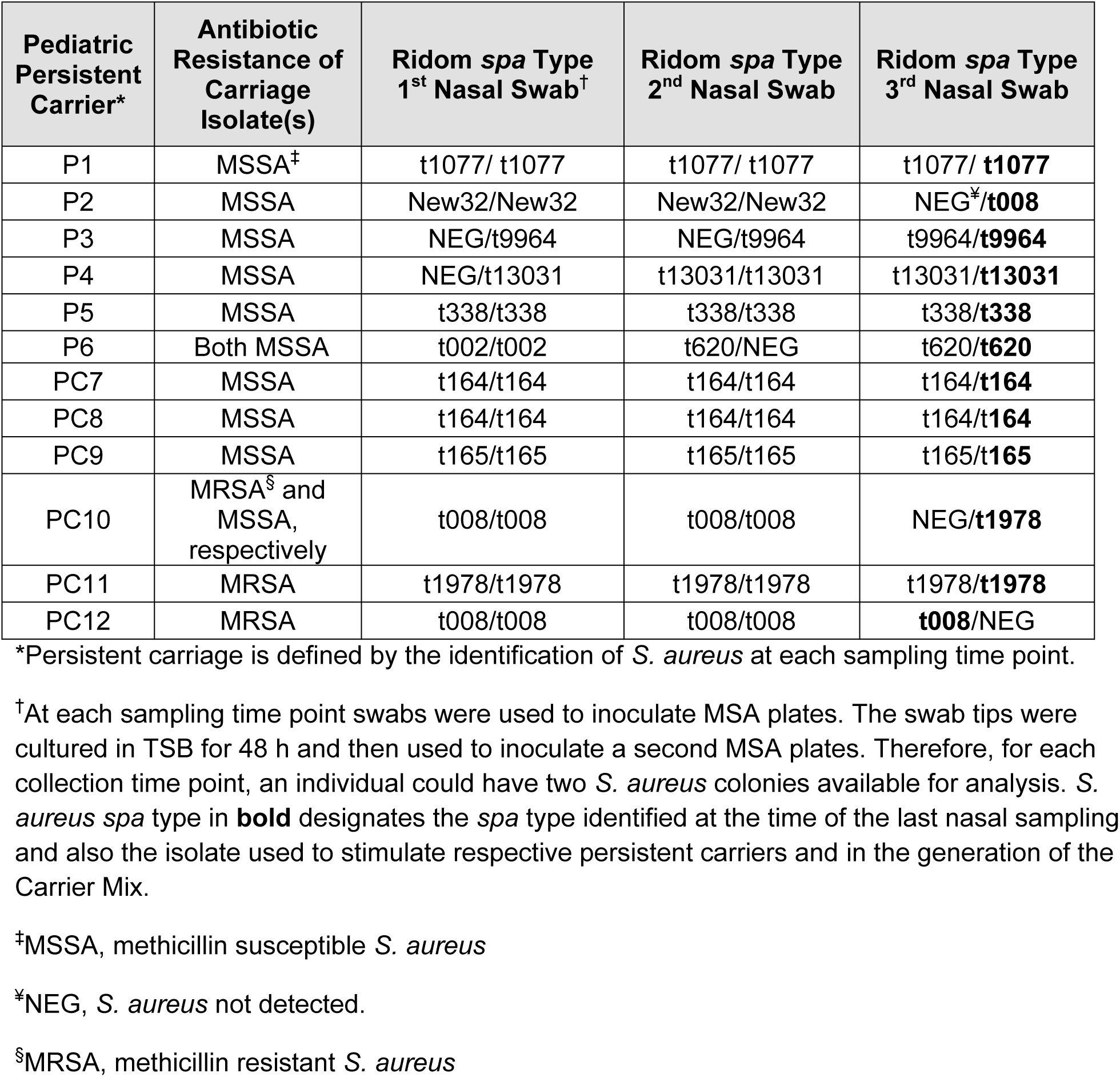
Strain characteristics of *S. aureus* strains isolated from pediatric persistent carriers.

### Technical validation of the targeted assay (TA)

Validation of selected genes was conducted by comparing individual genes using fold change values obtained from microarray data or the TA qPCR method at both module and individual gene levels. Eighty-four out of 90 genes correlated (defined by p<0.001) by comparing the microarray to the TA (Fig. 1 and Table S1). The genes listed on the x-axis of Fig. 1C correspond to the genes described in Table 3 and Tables S1 and S2. The module results obtained from the microarray, however, correlated well to those of the TA (Fig. 2), suggesting that the integrity of the module-based signature is more robust than that of individual genes. The module concept remains accurate both in relation to the data sets used to select the genes (33) and to the microarray data collected in the present report (Fig. 3). Based on these data, all genes tested remained in the TA even if on an individual level they did not correlate perfectly.

**Table 3.**
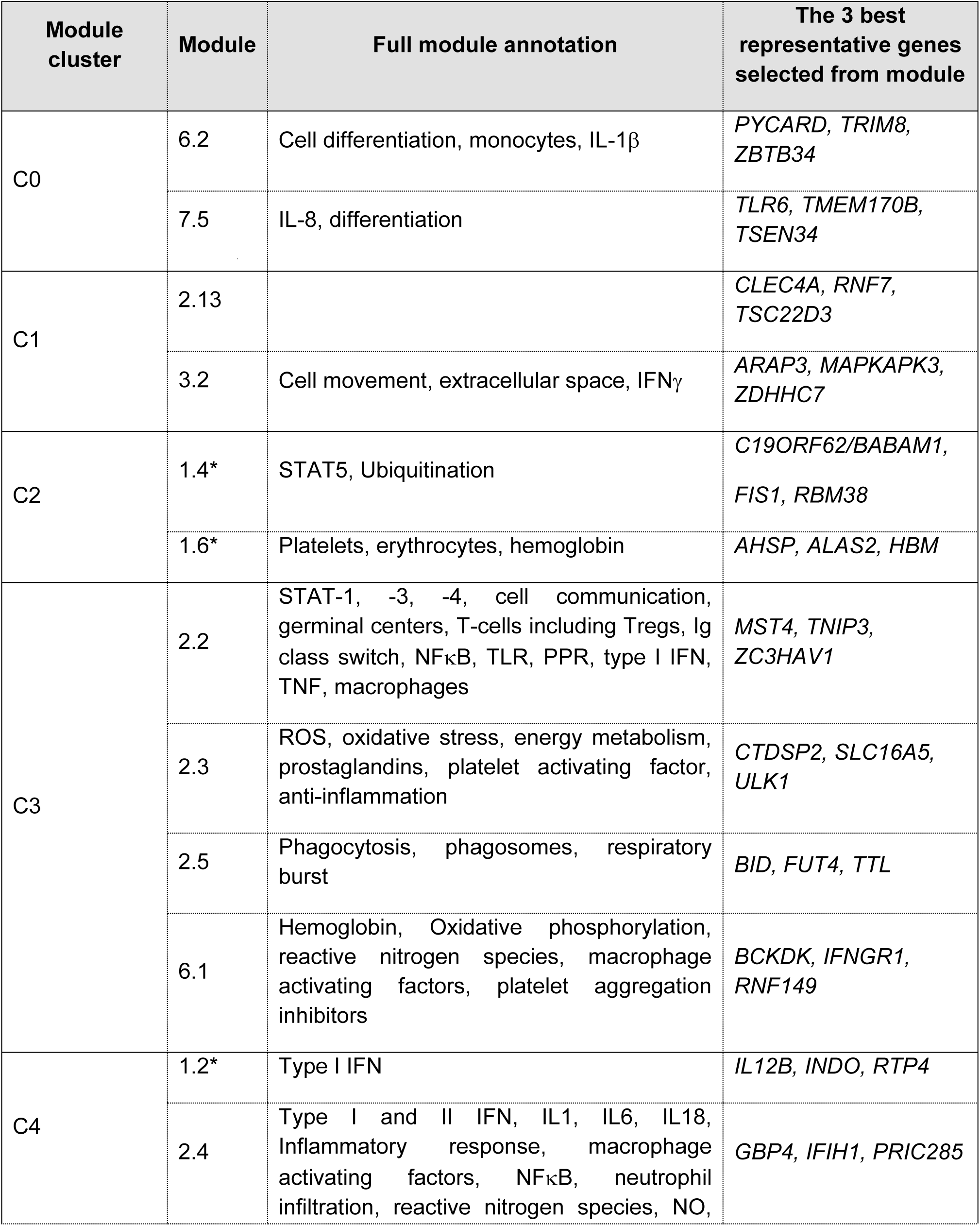
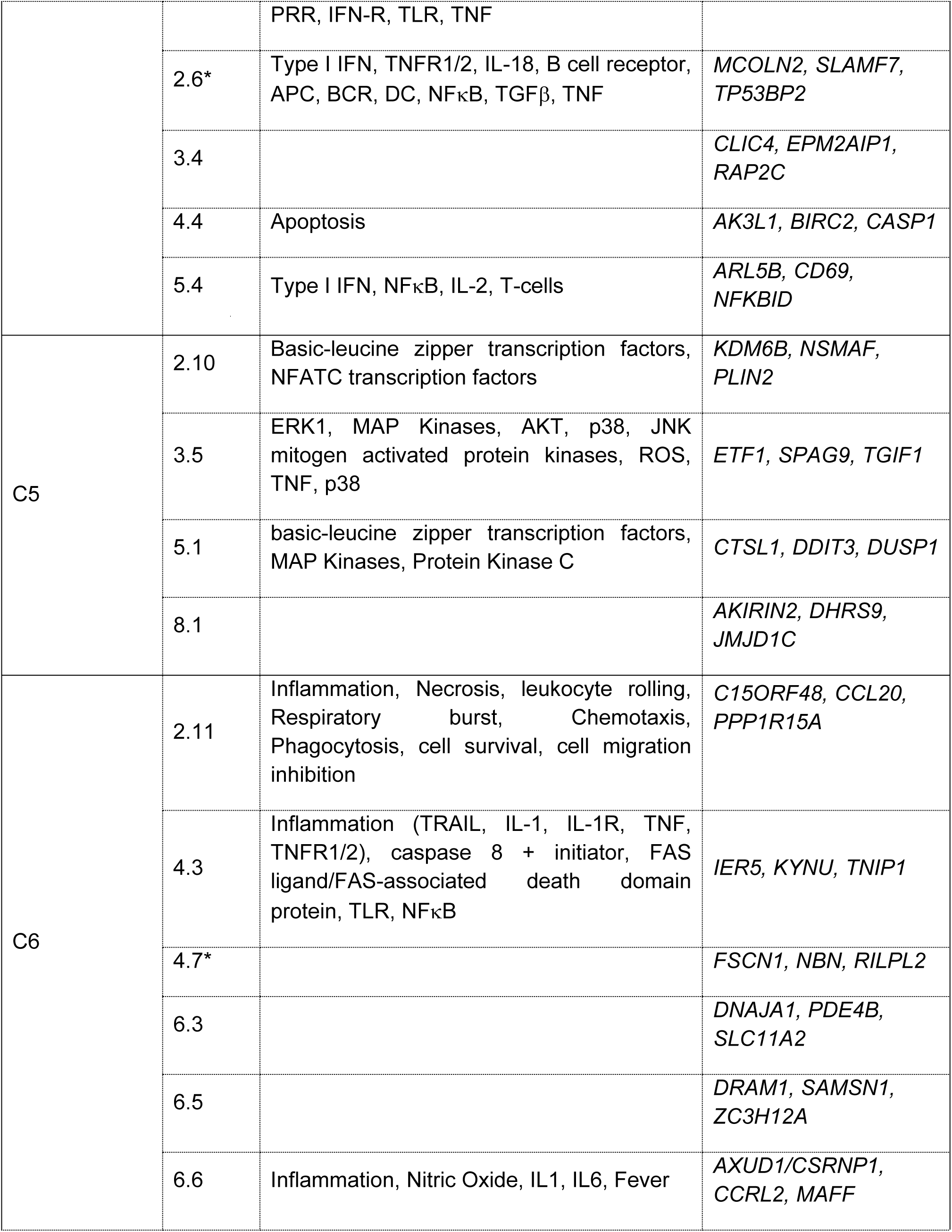
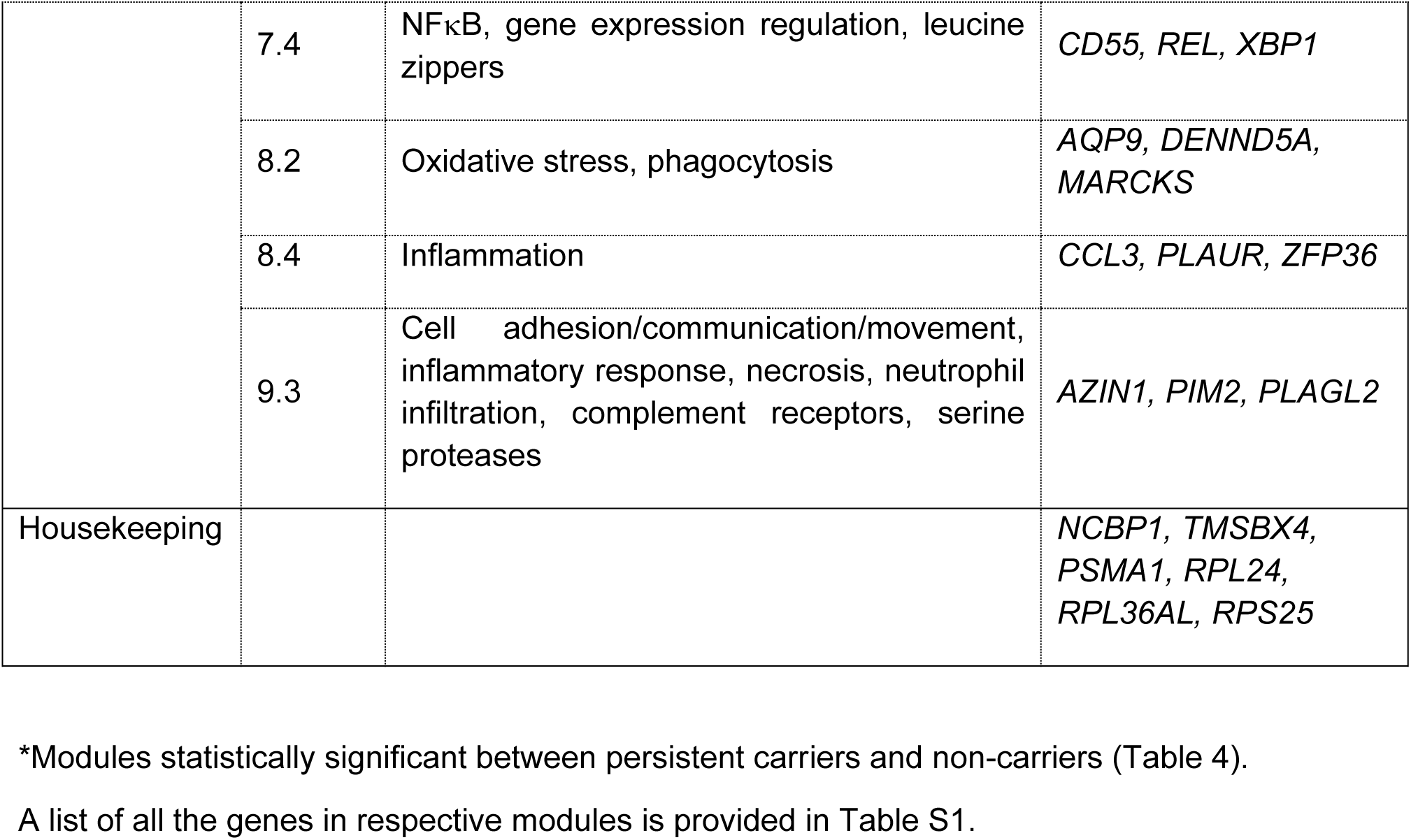
Modules with statistically significant expression differences between persistent and non-*S. aureus* carriers.

**FIG 1.**
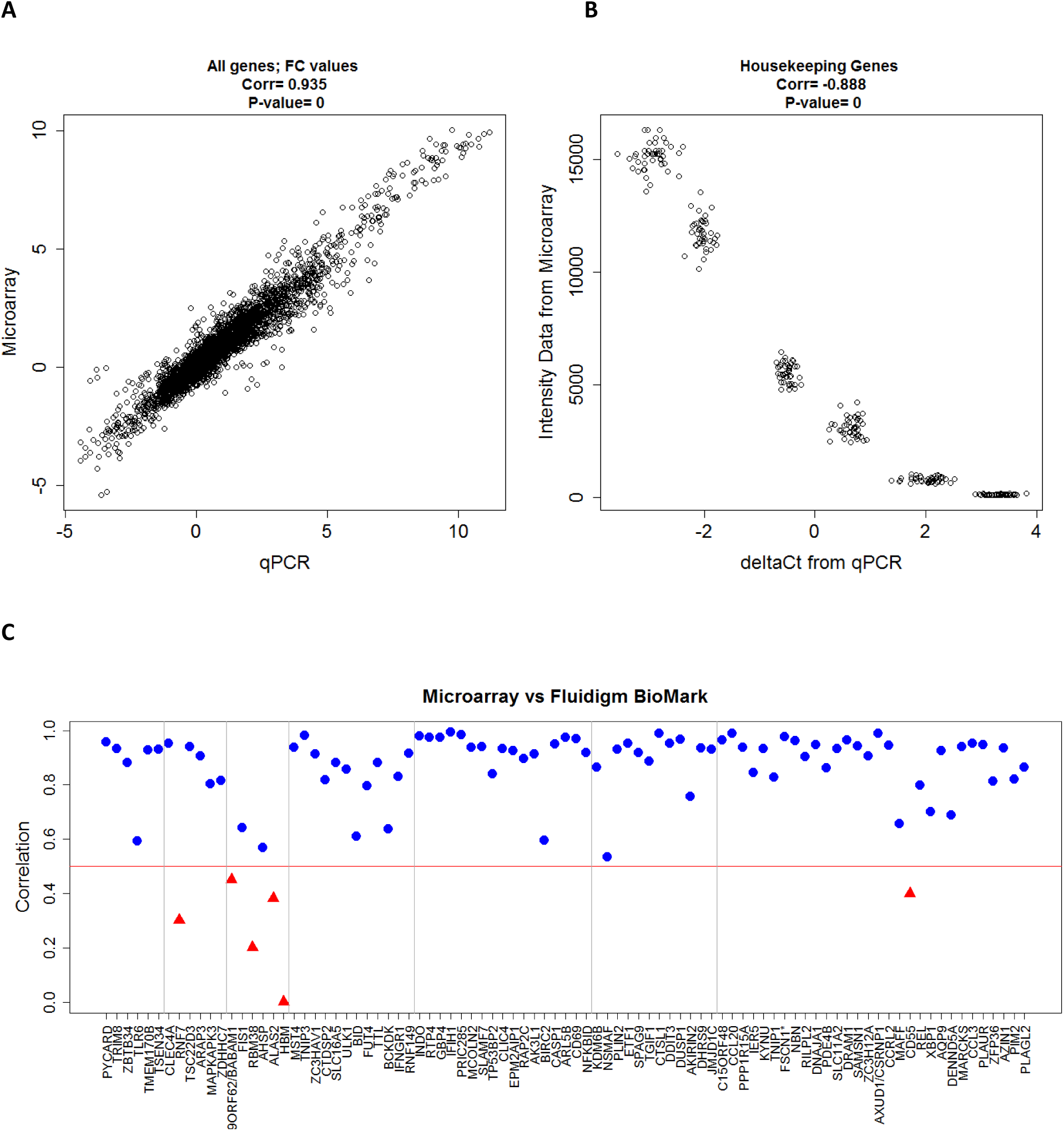
Spearman rank correlation scatter plots from genes represented in targeted and microarray assays. (**A)** Spearman Rank correlation values for each individual gene between microarray and the TA using log_2_ FC (fold change) values from 8 healthy adult donors. (**B)** Spearman Rank Correlation plot for selected housekeeping genes using intensity values from microarray and delta Ct values from the targeted assay from 8 healthy adult donors. The genes are, from left to right, *TMSB4X, RPS25, RPL24, RPL36AL, PSMA1*, and *NCBP1*. (**C)** Spearman Rank correlation values for each individual gene between microarray and TA using log_2_FC values. Values having a p-value less than 0.001 are represented by blue points, and values having a p-value greater than 0.001 are represented by red triangles. The red line indicates the 0.5 correlation level. The genes are ordered based on module, and the grey vertical lines separate the genes into the module cluster they represent.

**FIG 2.**
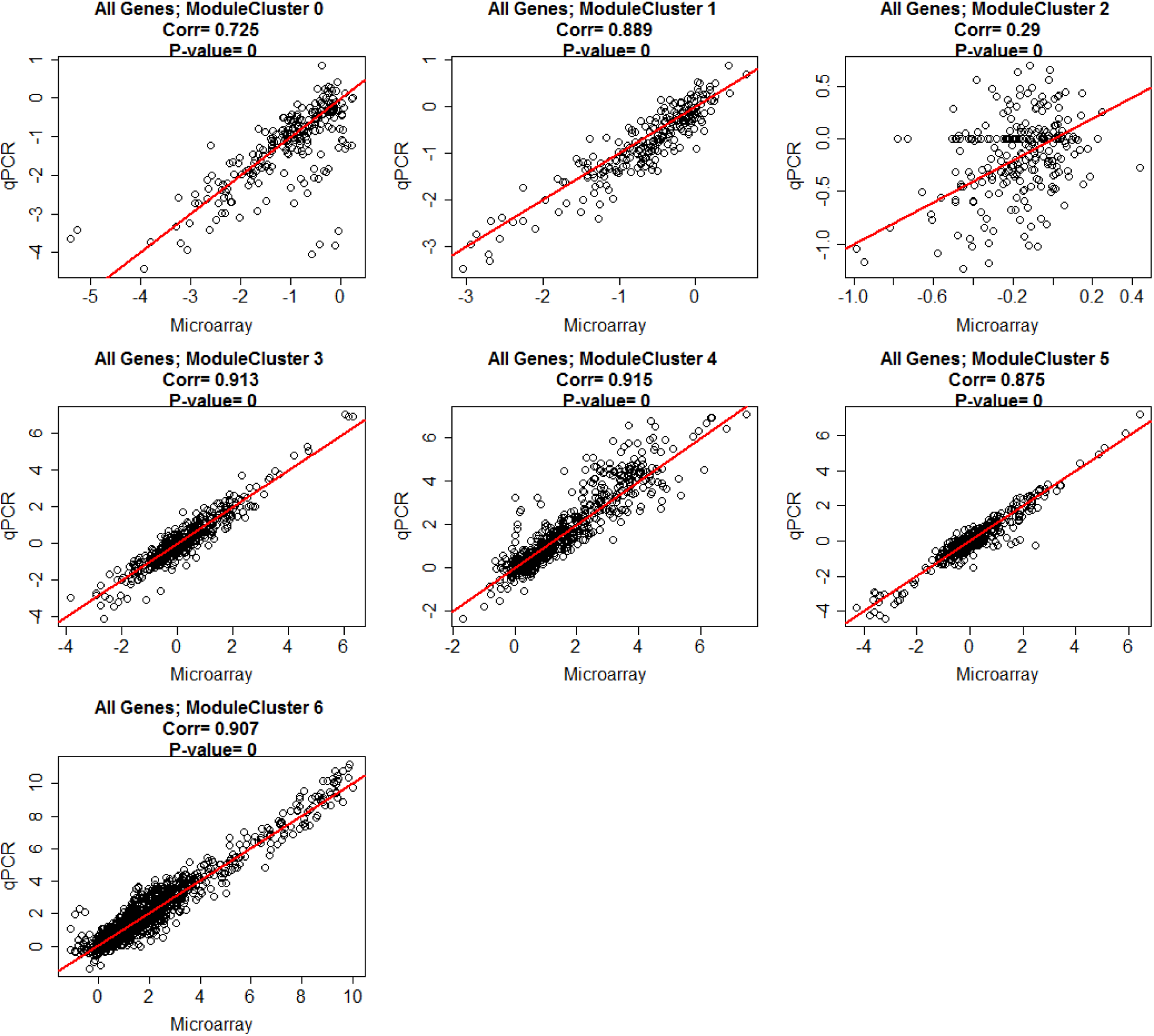
Spearman Rank correlation values for each individual gene in the different module clusters between microarray and TA using log_2_ transformed FC (fold change) values. Data is from 8 donors and 11 different stimulations in all plots, but the number of genes differ between the module clusters (Module Cluster 0: 6 genes, Module Cluster 1: 6 genes, Module Cluster 2: 6 genes, Module Cluster 3: 12 genes, Module Cluster 4: 18 genes, Module Cluster 5: 12 genes, Module Cluster 6: 30 genes). [Total number of data points per plot: Module Cluster 0: genes n = 528 (8 donors × 11 stimulations × 6 genes), Module Cluster 1: n = 528 (8 donors × 11 stimulations × 6 genes), Module Cluster 2: n = 528 (8 donors × 11 stimulations × 6 genes), Module Cluster 3: n = 1056 (8 donors × 11 stimulations × 12 genes), Module Cluster 4: n = 1584 (8 donors × 11 stimulations × 18 genes), Module Cluster 5: n = 1056 (8 donors × 11 stimulations × 12 genes), Module Cluster 6: n = 2640 (8 donors × 11 stimulations × 30 genes)]

**FIG 3.**
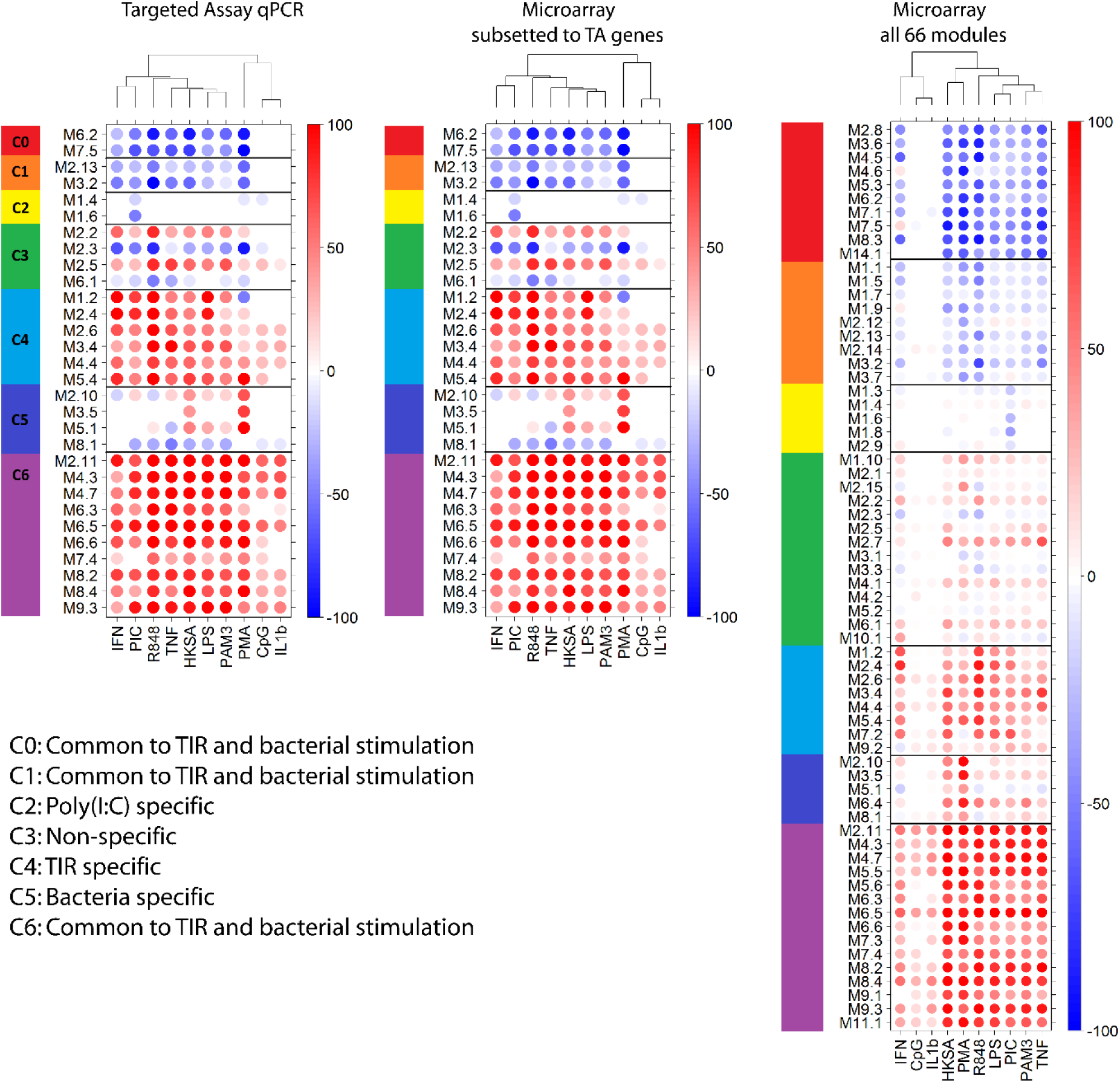
Module responses from TA (30 modules), microarray subset to TA genes and modules, and the full modular transcriptome repertoire (66 modules) from the full microarray from the 8 healthy adults used for method development. The module responses between TA and microarray subset to TA genes are identical. The full 66 modules show the same characteristics expected from (33): module cluster 2 is poly (I:C) specific, module cluster 5 is bacterial specific, module cluster 3 is non-specific, and the remaining clusters are TIR and bacterial specific. The module response pattern seen in the full 66 modules can be seen in both TA and the microarray subset.

### The effect of different sample sources on the TA

TA robustness to sample source was tested by comparing RNA analyses obtained from fresh whole blood and PBMCs and to previously frozen PBMCs, all sourced from the same donors and blood draws. The correlations between the different sources show that at 6 h post-stimulation the 3 distinct RNA sources correlate reasonably well (Fig. 4A), whereas at 2 h post-stimulation frozen PBMCs do not correlate well with either whole blood or freshly isolated PBMCs (Fig. 4A) suggesting that 2 h may not be optimal for conducting the TA when working with previously frozen PBMCs. Heat maps of module responses for each stimulation show similar patterns across RNA sources at 6 h but not at 2 h (Fig. 4B). These data suggest that previously frozen PBMCs respond indistinguishably from freshly isolated PBMCs or fresh whole blood in the TA if longer activation times are tested, thereby providing a tool that makes it possible to obtain accurate module information for fresh and frozen PBMCs even though the original modules were developed for the analysis of RNA derived from whole blood.

**FIG 4.**
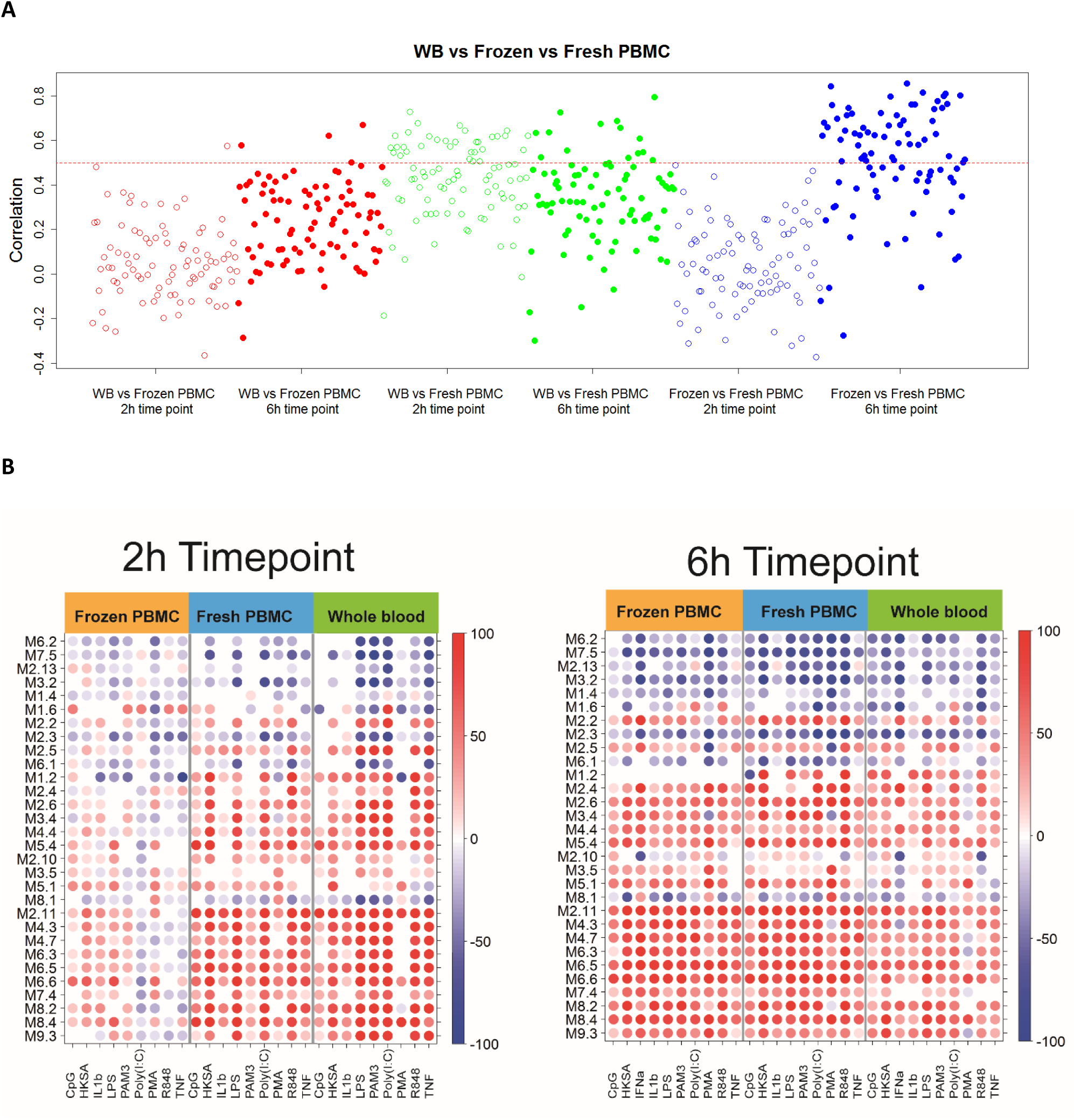
Comparisons between whole blood (WB), freshly isolated PBMCs, and previously frozen PBMCs from 4 healthy adult donors. (**A)** Correlation values (of log_2_FC to non-stimulated self) comparing WB, fresh PBMCs, and frozen PBMCs. Open circles represent the 2 h time point and closed circles represent 6 h time point. The dashed red line indicates the 0.5 correlation value. At the 2 h time point frozen PBMCs show low correlation to fresh PBMCs or WB. The correlation improves at 6 h. Fresh and frozen PBMCs are most similar followed by a similar correlation pattern for WB to fresh PBMC and WB to frozen PBMCs. **(B)** Module plot. The figure represents the modular activity of each donor for the 30 TA modules. Spot intensity indicates changes in transcript abundance from baseline. No clustering was applied to either samples or Modules. Intensity for each module represents the percent of module probes that are up (red) or down (blue) regulated in a sample. At the 2 h time point frozen PBMCs differ from fresh PBMCs and WB, but at the 6 h time point this difference is no longer detectable.

### PBMC responses to different stimuli

We used the TA to assess gene expression profiles associated with innate immunity by stimulating PBMCs harvested from pediatric persistent or non-carriers of *S. aureus* with various stimuli.

Although the responses observed to the various stimuli differed, responses to respective stimuli were similar between groups (Fig. 5, Fig. S1). Differences in responses observed between persistent carriers and non-carriers were observed for M7.5 following PAM3 stimulation (p=0.04, Fig. 5), M2.5 following stimulation with the Carrier Mix (p=0.01, Fig. 5), and M2.13 following CpG stimulation (p=0.004, Fig. S1). Some modules showed large donor-to-donor variation (*e.g.*, M1.6 showed donor to donor variation for all stimulations) while other module responses were relatively stable across donors (*e.g*., M5.4 and M8.2 in the Carrier Mix panel). M8.2 is down-regulated across all donors after Carrier Mix stimulation but up-regulated after PAM3 stimulation (Figure 5). The other stimulation profiles showed donor to donor variation. For example, M5.4 is up-regulated in all donors after Carrier Mix stimulation, but only in a few donors following stimulation with PAM3 or other stimulations, and responses in M1.2 are variable in all donors following stimulation with either Carrier Mix (Fig. 5), TNF, or CpG, whereas M1.2 responses are up-regulated following stimulation with either PAM3, IL-1β, or HKSA (Fig. S1).

**FIG 5.**
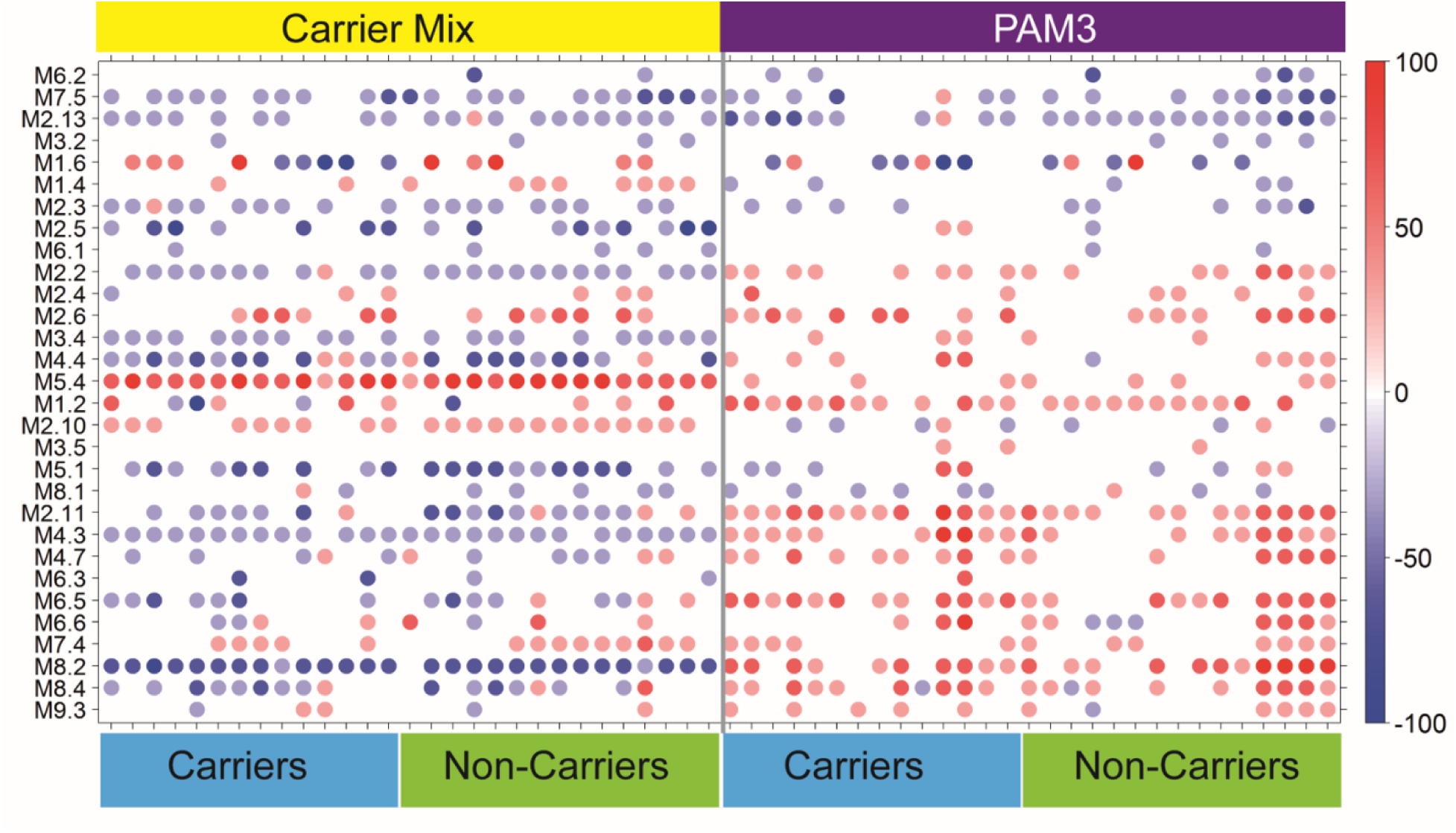
Responses to a 6 h stimulation with either a mix of all clinical carrier strains (Carrier Mix) or PAM3 are similar between S. aureus carrier status groups. The module plot represents the modular activity of each donor for the 30 TA modules. Spot intensity indicates changes in transcript abundance from baseline. No clustering was applied to either samples or modules. Intensity for each module represents the percent of module probes that are up (red) or down (blue) regulated in respective samples. The modular responses are different between the mix of all clinical carrier strains and PAM3, but no differences were found between carriers and non-carriers. Some modules show larger donor-to-donor variations (*e.g.*, M1.6) while others are relatively stable across donors (*e.g.*, M5.4 and M8.2 in the Carrier Mix panel).

The responses to the Carrier Strain and the Carrier Mix were, although not statistically significant, slightly different both at a gene level (Fig. 6A) and the module level (Fig. 6B). Some modules showed large donor-to-donor variation (*e.g*., M1.6, and M1.2) while others were relatively stable across donors (*e.g*., M5.4, M4.3, and M8.2) pointing to the relative importance of these modules following stimulation with bacterial antigens. When looking at the module responses in an individual stimulated with the Carrier Mix compared to the response elicited following stimulation with their respective carrier strain, small and subtle differences were observed (Fig. 6B). Although not statistically significant, these differences still may contribute to the understanding persistent carriage of specific *S. aureus* isolates in these individuals.

**FIG 6.**
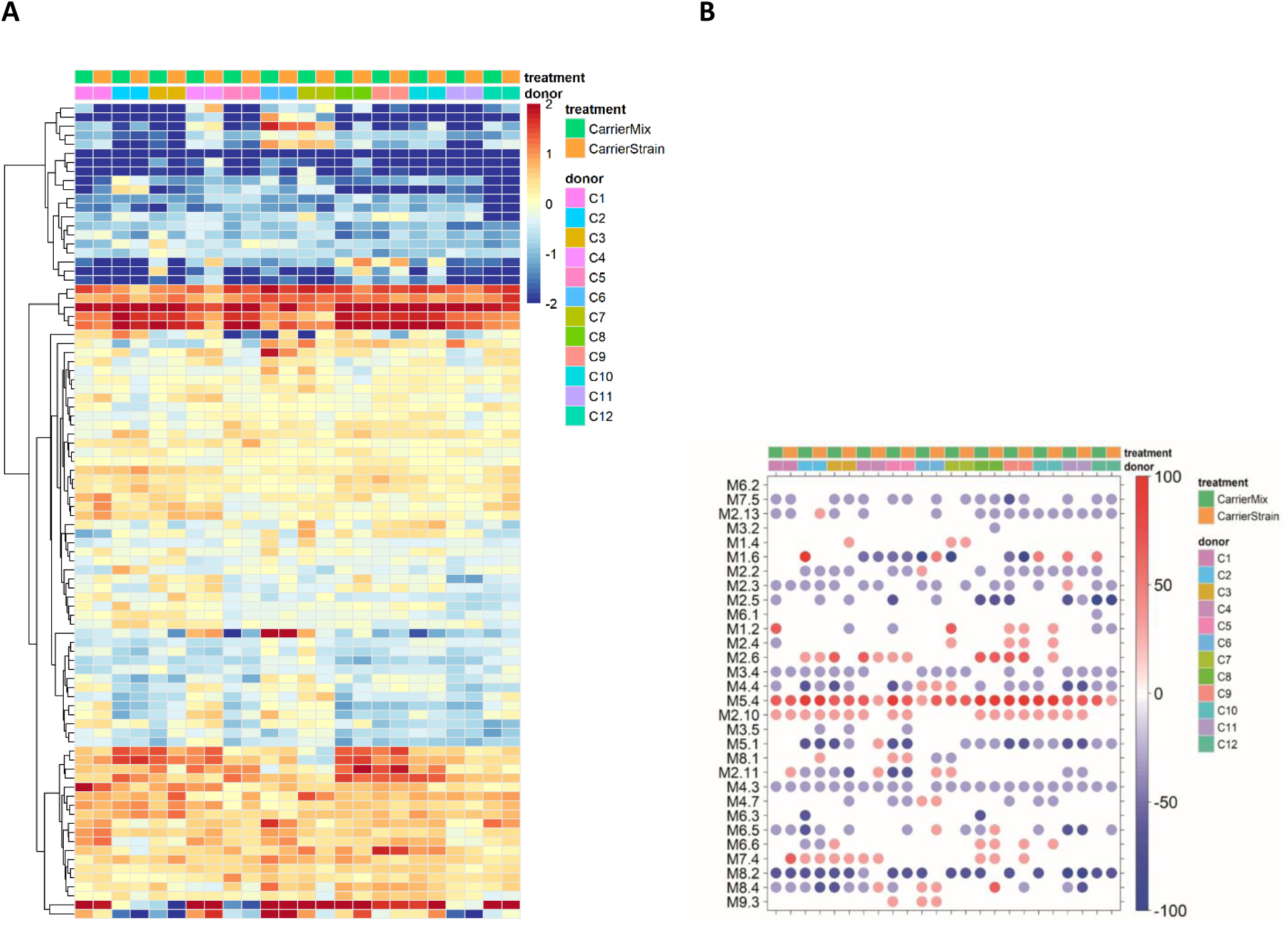
Responses by PBMCs isolated by persistent carriers to donor specific clinical carrier strains or the mix of all clinical carrier strains following stimulation for 6 h. **(A)** Heat map comparing gene expression profiles after stimulation with either the donor specific clinical carrier strain or a mix of all clinical carrier strains. Heat maps represent a hierarchical clustering of transcripts responding to each stimulus (compared to non-stimulated self). Changes versus the non-stimulated condition are represented by a color scale: red = up regulated; blue = down regulated; yellow = no change. Some donors show more variability than others. **(B)** The module plot represents the modular activity of each donor for the 30 TA modules. Spot intensity indicates changes in transcript abundance from baseline. No clustering was applied to either samples or modules. Intensity for each module represents the percent of module probes that are up (red) or down (blue) regulated in a sample. Some modules show larger donor-to-donor variation (*e.g*., M1.6) while others are relatively stable across donors (*e.g*., M5.4 and M8.2) suggesting the relative importance of these modules in bacterial responses.

### PBMC responses to live or dead bacteria

Clear differences were observed in responses elicited following stimulation with heat-killed *S. aureus* compared to stimulation with live carriage isolates (Fig. 7 and Table 4). Modules showing statistically significant differences in gene expression profiles following stimulation with either heat-killed or live bacteria were annotated as “Type I IFN” (M1.2, M2.2, M2.6, M5.4), “Inflammation” (M4.3, M8.4, M2.11), “Apoptosis” (M4.4), “Transcriptional Regulation” (M7.4, M5.1, M2.10, M1.4), “Phagocytosis” (M2.5, M8.2), or none or poorly defined annotations (M2.3, M2.13, M3.4, M4.7, M6.5, M7.5). Only 9 of 30 modules were not different following stimulation with either live or heat-killed bacteria and 16 modules showed a consistent pattern in both carriers and non-carriers. In addition, seven modules (M1.2, M1.4, M1.6, M2.6, M2.10, M4.7, and M5.1) showed small differences to stimulation with live bacteria between carriers and non-carriers (Fig. 7). Especially interesting are the small differences observed in the expression profiles observed for modules M1.4 and M1.6 (the two modules from the poly [I:C] specific cluster) that were up-regulated in non-carriers stimulated with live bacteria, but down-regulated after heat-killed bacteria stimulation. The expression profiles for these same modules were unchanged when PBMCs harvested from persistent carriers were stimulated with either live or dead bacteria (Fig. 7). Recently, induction of IFNβ via a TIR-domain-containing adaptor protein (TRIF)-dependent pathway (34) following stimulation with live pathogenic bacteria was identified as a potential mechanism for distinguishing between stimulation with either live or dead organisms or between pathogenic or commensal bacteria (34). This suggests that individuals susceptible to persistent carriage with *S. aureus* may possess differences in their live/dead bacteria recognition pathway.

**Table 4.**
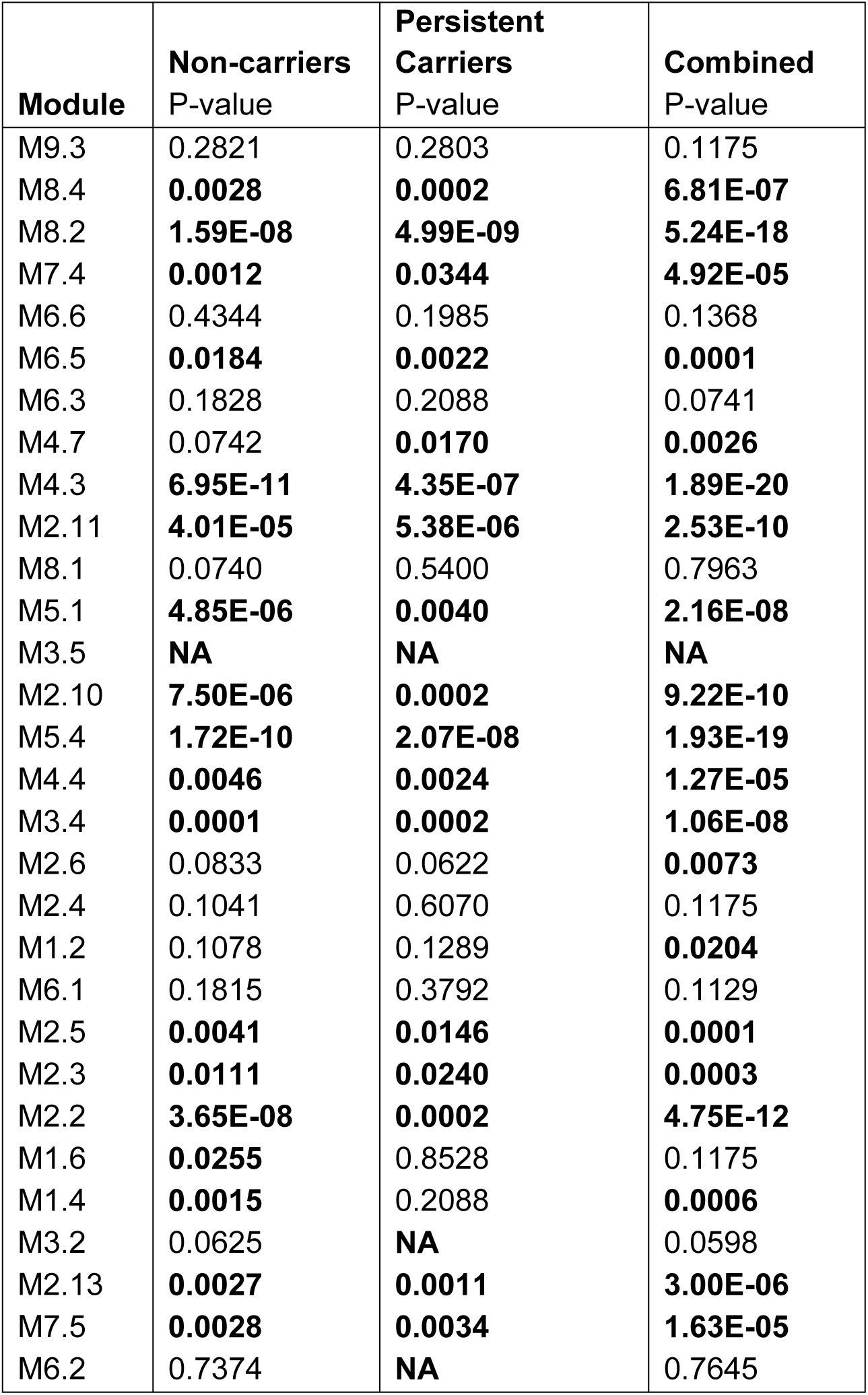
Differences in module scores between heat-killed and live bacteria. FDR corrected P-values for each module. P-values in bold are <0.05. NA indicates that the tested values were identical and the t-test did not return a significant p-value.

In order to obtain a better understanding of the different responses observed following stimulation with live versus dead bacteria between persistent carriers and non-carriers, we performed RNAseq on unstimulated cells, cells stimulated with heat-killed bacteria, and cells stimulated with live bacteria from a subset of the donors representing the average response of the respective groups (Fig. S2). A principal component analysis (PCA) shows that samples stimulated with live bacteria are different from those stimulated with dead bacteria or left unstimulated (Fig. 8A). Furthermore, differential expression of genes (Table 5 and Table S3) and modules (Fig. 8B) is greater following stimulation with live bacteria compared to stimulation with heat-killed bacteria. Furthermore, non-carriers and persistent carriers differ in their module responses and differences between persistent carriers and non-carriers detected by the targeted assay are more pronounced when using RNAseq (Fig. 8C).

**Table 5.** Differentially expressed genes from RNAseq analysis.

| Donor group | Comparison | FDR<0.05 | Top 5 Biological functions from Pathway Studio |
| --- | --- | --- | --- |
| Persistent Carrier | Live bacteria vs. NS | 5171 | Microtubule Cytoskeleton<br>Centriole Duplication and Separation<br>P2RXs -> Synaptic Transmission<br>Kynurenine Metabolites in Neurotoxicity and Neuroprotection<br>SWI/SNF BRG1/BAF in Chromatin Remodeling |
| Persistent Carrier | HKSA vs. NS | 101 | mRNA Transcription and Processing<br>Epinephrine/Norepinephrine<br>SWI/SNF BRG1/PBAF in Chromatin Remodeling<br>Mast-Cell Activation without Degranulation through CYSLTR1/CYSLTR2 Signaling<br>Histone Methylation |
| Non-carrier | Live bacteria vs. NS | 6535 | Golgi to Endosome Transport<br>SRCAP in Chromatin Remodeling<br>Actin Cytoskeleton<br>PTGER2/3 -> Inflammation-Related Expression Targets<br>Synaptic Potentiation by PKMZ in LTP Maintenance - Basal State |
| Non-carrier | HKSA vs. NS | 20 | Telomere Maintenance<br>Single-Strand Nucleotide Excision<br>Tight Junction Assembly (JAMs)<br>Neutrophil Recruitment and Priming<br>SIRT2 Signaling in Aging |

**FIG 7.**
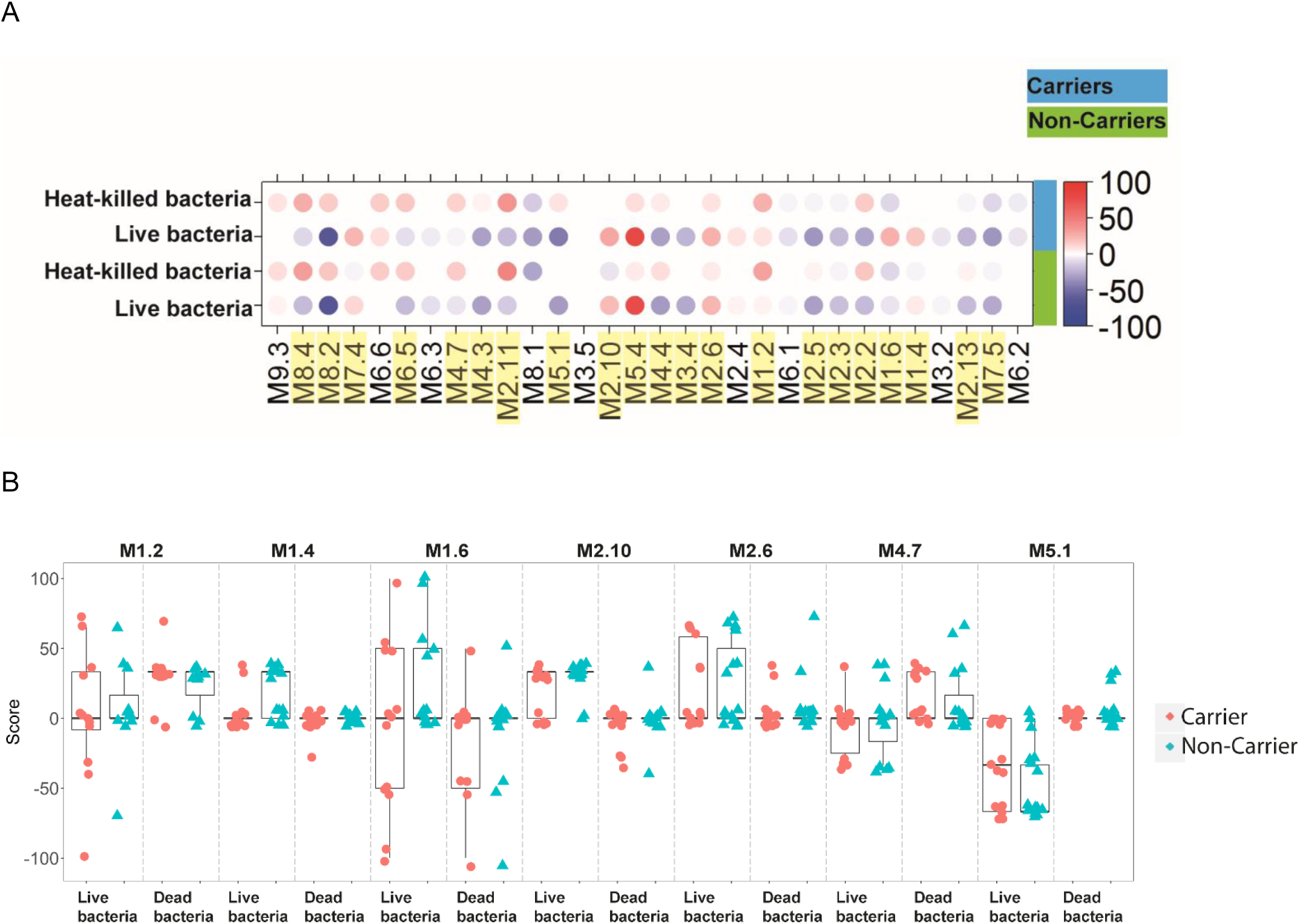
Responses to live verses dead bacteria. **(A)** The module plot represents the mean modular activity of each donor group for the 30 TA modules. Spot intensity indicates changes in transcript abundance from baseline. No clustering was applied to either samples or modules. Intensity for each module represents the percent of module probes that are up (red) or down (blue) regulated in a sample. Yellow highlights denote modules where the module score is statistically significantly different between live and dead bacteria in at least one of carriers and non-carriers, or both groups combined **(B)** The individual module score for the 7 modules where Carriers showed a different pattern in module scores than seen in Non-carriers.

**FIG 8.**
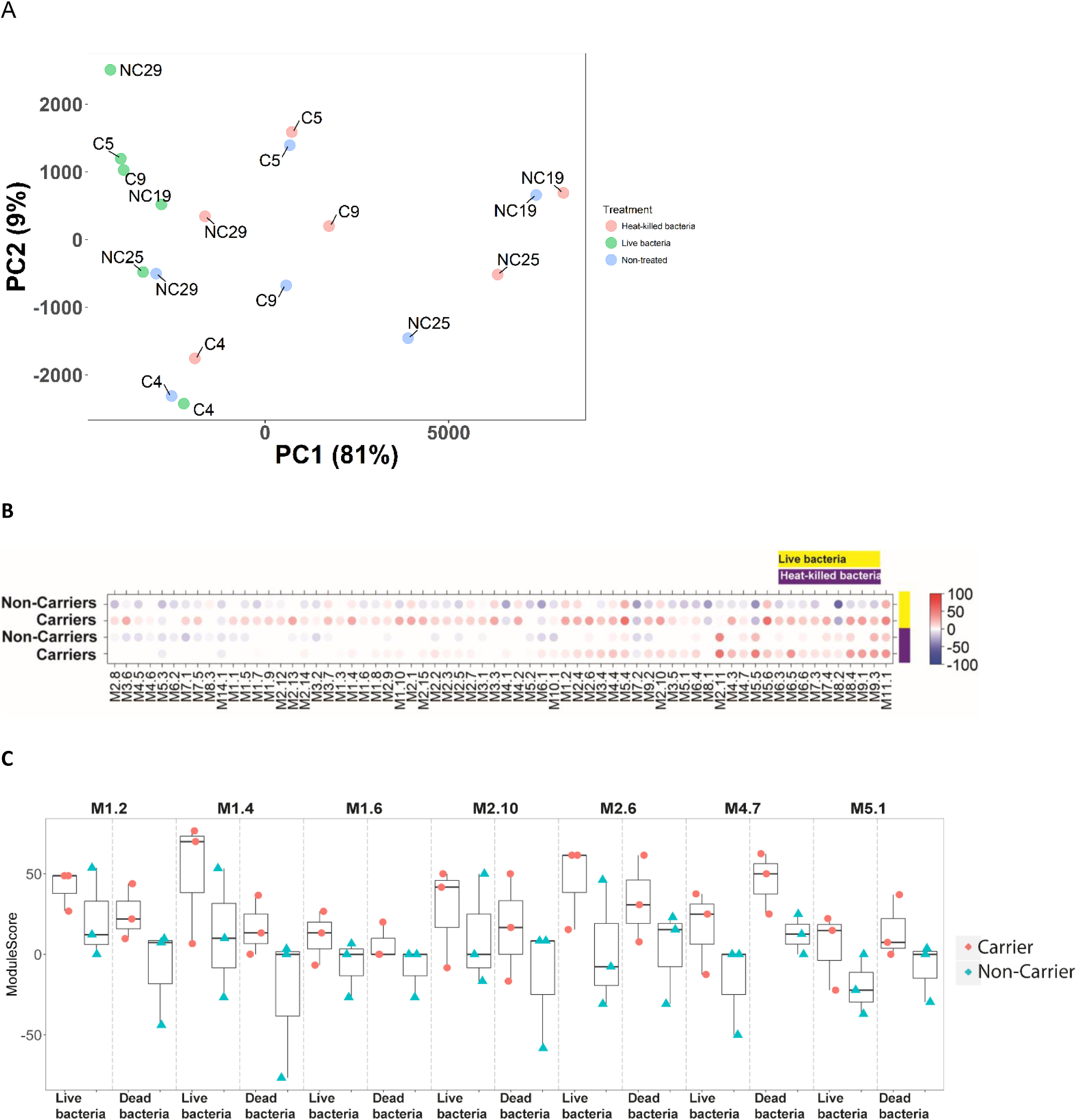
Principle Component Analysis plot and Module plot. **(A)** Principal Component Analysis (PCA) plot showing variability in the data. Treatments are indicated by different colors: blue = NS [not stimulated], red = Heat-killed bacteria, green = Live bacteria. **(B)** Mean modular activity of each donor group for the 30 TA modules. Spot intensity indicates changes in transcript abundance from baseline. No clustering was applied to either samples or modules. Intensity for each module represents the percent of module probes that are up (red) or down (blue) regulated in a sample. **(C)** The individual module scores for the modules identified in Figure 7A-B.

We were next interested in identifying genes differentially activated in carriers and non-carriers. Because we hypothesized that differences would be minor due to our small data set, we tried two different approaches aimed at identifying potential differences: (1) statistical testing using a general linear model using negative binomial distribution with no random effects (edgeR) and (2) measuring the “deviation from the perfect line” (the dot product). Any deviation from this line could be due to a difference in responses to live bacteria in the different donor groups. When applying these approaches to our data, 20 genes were identified as different between carriers and non-carriers when using the statistical model (Table 6) and a total of 367 genes showed an altered profile for persistent carriers compared to non-carriers using this “deviation from the perfect line” approach (Fig. 9 and Table S4). The overlap between the two different gene lists were rather low, only 5 genes were found with both methods (Table S4), but the remaining genes were all close to the cutoff used for the “deviation from the perfect line”.

**Table 6.**
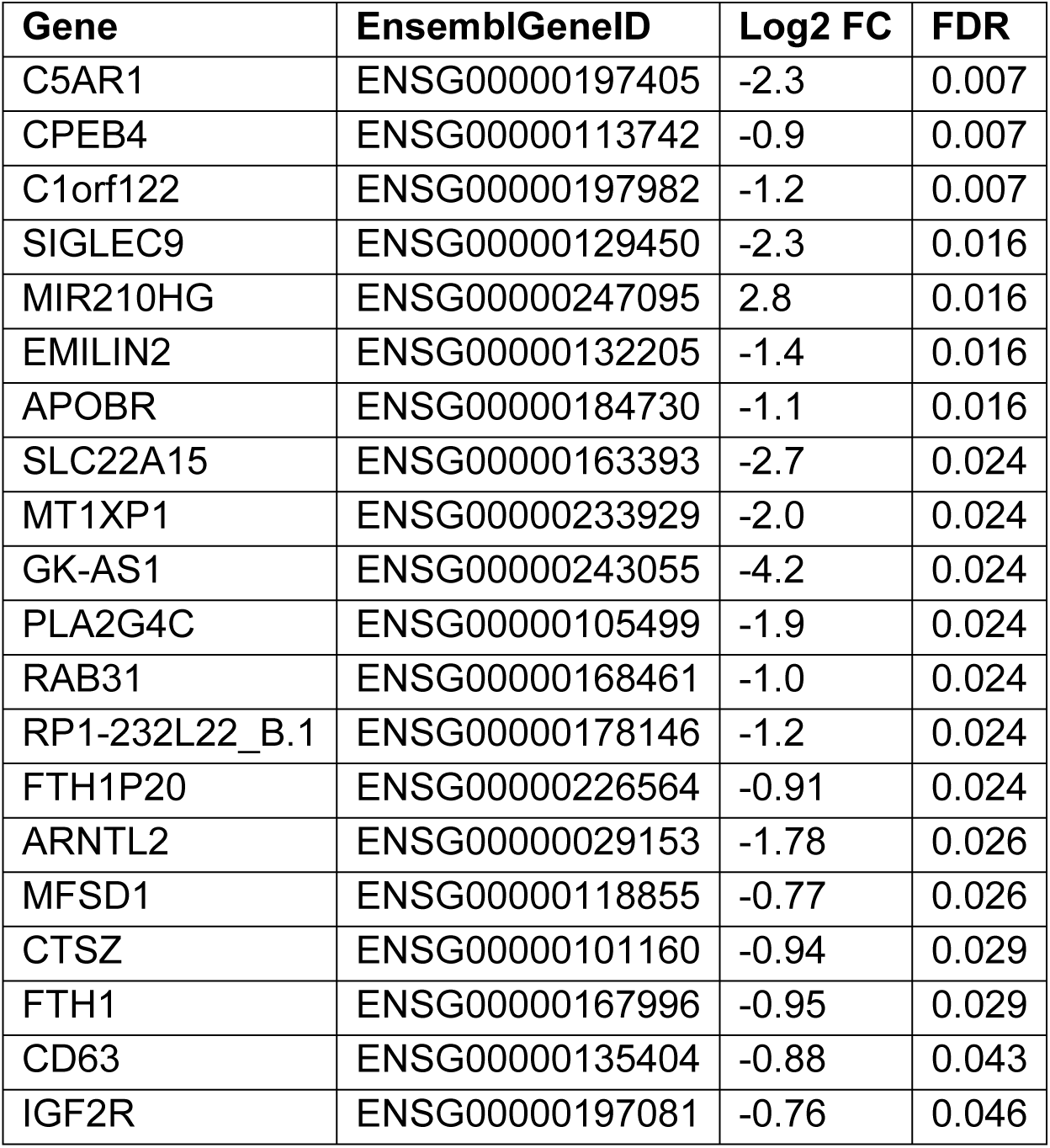
Differentially expressed genes for live vs. dead *S. aureus* between Carriers and Non-Carriers

**FIG 9.**
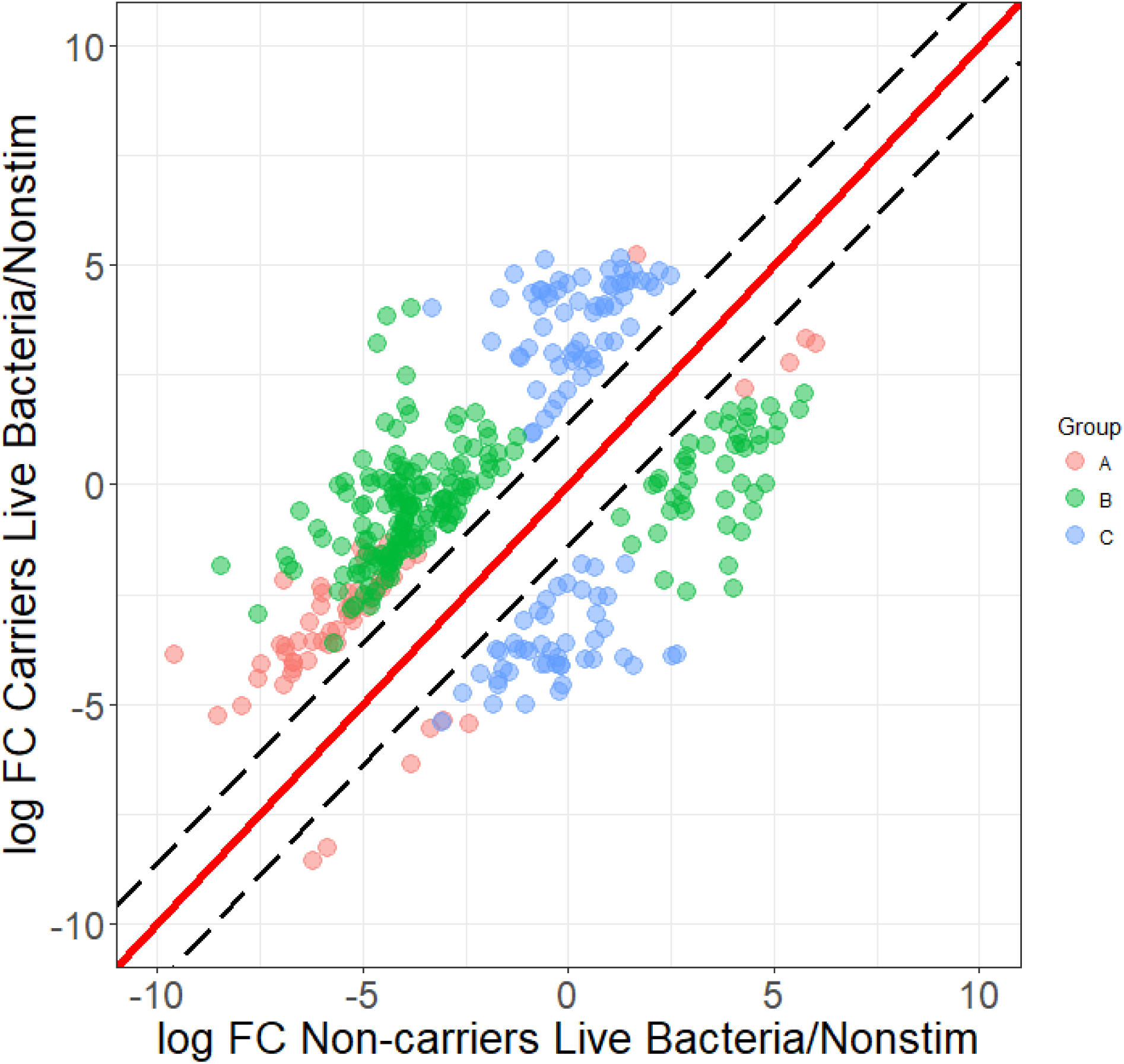
Defining the variation between carrier and non-carrier gene signatures after co-culture with live bacteria. Black dashed lines indicate the interval assumed to be normal variation (±2 SD). All points indicate genes outside the normal variation and with an FDR-corrected p-value <0.05 in at least one donor group (live bacteria vs. non-stimulated) (n = 367). Three groups of genes can be identified based on the cutoffs they pass: A: 2*SD and FDR<0.05 for both non-carriers and carriers (red points, n = 60), B: 2*SD and FDR<0.05 for non-carriers but not persistent carriers (green points, n= 198), C: 2*SD and FDR<0.05 for persistent carriers but not non-carriers (blue points, n = 109).

## Discussion

Interrogation of the human genome can be carried out by either conducting analyses that define either its architecture or measure its output. Genome-wide association studies represent an example of the former approach designed to identify genes or pathways that are determined by heredity and do not change over time that may be associated with various immune-related, chronic, and infectious diseases (35–40). By contrast, genetic approaches that measure transcript abundance *e.g*. the TA, are largely affected by environmental stimuli such as infection/colonization status.

The modular transcriptional repertoire analysis pioneered by our laboratories was used as a basis for the development of the TA and used to identify differences in the transcriptome between pediatric individuals determined to be either persistent nasal carriers or nasal non-carriers of *S. aureus* (29, 33, 41, 42). The module concept is based on co-expression of genes that may or may not be related and individual genes represented in respective modules may not necessarily be directly involved with the function of the modules. This assay is meant to be analyzed and interpreted at the modular level where individual genes comprising each module contribute to a profile that can be used to understand potential defects in innate immune responsiveness, not as a tool to pinpoint the exact gene responsible for respective phenotypes.

The modules used in the present study were specifically developed for analyses of stimulated whole blood (33), but the underlying framework is the same as what has been previously described (41). More importantly, the TA described in the present report allowed for frozen PBMCs to be analyzed, producing results indistinguishable from similarly treated fresh whole blood. Also, the time and cost reduction that comes by using the TA makes it a very good first screening tool, especially when there are many samples to process.

The advantage the modular approach which the TA described here is based on is highlighted by prior work that identified a type I interferon signature in association with systemic lupus erythematosus not previously identified using traditional approaches (43–45) that resulted in the development of anti-INF-α monoclonal antibody-based therapies for the treatment of this disease (46). In the context of infectious diseases, comparison of principal gene-expression patterns identified a role for neutrophils (previously believed to be uninvolved in protection against *Plasmodium* infections) in resistance to malaria (47) and the module approach was successfully used to define transcriptome signatures of infections associated with a variety of pathogens and that may be developed as a surveillance tool designed to monitor disease progression or reactivation (44).

An understanding of the immune components that coordinate the elicitation of protective immunity to respective pathogens is requisite to the development of therapeutics and vaccines. This is the primary reason why effective vaccines have been developed for the prevention of diseases resulting from infections with *Streptococcus pneumoniae*, *Haemophilus influenzae* type B, *Neisseria meningitidis*, and toxigenic bacteria causing diphtheria, pertussis and tetanus but not for organisms such as *S. aureus*, numerous other bacteria, protozoans, or parasitic helminths. Gene-expression analyses provide an opportunity for the identification of genes/pathways that may play roles in defining *S. aureus* carriage phenotypes that can used to better understand the nature of immune response profiles associated with susceptibility to *S. aureus* carriage and/or infection.

This study is different from other infectious disease gene expression analyses studies in that the groups studied were defined by their respective *S. aureus* carriage phenotype and not by infection status. Specifically, we assessed differences in gene expression profiles between persistent and non-carriers of *S. aureus* since these two carriage phenotypes are the most biologically distinct (18, 19). Furthermore, some investigators have suggested that because immune response profiles of intermittent carriers and non-carriers to *S. aureus* are more similar they should be considered a single carriage phenotype (18). For example, persistent carriers are at higher risk of autoinfection despite having significantly higher specific IgG and IgA titers to *S. aureus* virulence factors than non-carriers and intermittent carriers and artificially inoculated *S. aureus* mixtures survived significantly longer in persistent carriers than they did in intermittent and non-carriers (18, 48). Furthermore, decolonized persistent carriers inoculated with a mixture of *S. aureus* isolates were more likely to be recolonized with their original resident strain (18, 19). Based on these basic physiologic/immunologic differences between persistent carriers and intermittent/non-carriers we hypothesized that the gene expression profiles between these two groups following PBMC activation with various stimuli would also differ.

Gene expression profiles that differed between persistent carriers and non-carriers were observed in M2.13 (cytoskeleton, cell cycle), M2.5 (phagocytosis, phagosomes, respiratory burst), and M7.5 (IL-8, differentiation) following stimulation with CpG, Carrier Mix, and PAM3, respectively. When PBMCs isolated from persistent carriers were stimulated with either their respective live carrier strains or the Carrier Mix differences in expression profiles, although not statistically significant, differed both at the gene and module levels supporting prior observations regarding the intimate association between persistent carriers and their respective persistent carriage isolates.

Differences in gene expression profiles were also observed between persistent carriers and non-carriers following stimulation with either heat-killed or live bacteria. A change in gene expression profiles of non-carriers was observed in modules associated with type I interferons, inflammation, apoptosis, transcriptional regulation or phagocytosis (M1.4, M1.6) when stimulated with live bacteria but not with dead bacteria. Surprisingly, there was little change in the expression profile observed for persistent carriers when stimulated with either live or dead bacteria. Additional RNAseq analyses were then conducted to uncover additional differences in responses to live/dead bacteria between persistent and non-carriers by comparing unstimulated cells, cells stimulated with heat-killed bacteria, and with live bacteria on a subset of participants. This analysis also uncovered differences in gene expression profiles between persistent carriers and non-carriers. Specifically, unique process networks for non-carriers were related to IL-1 and IL-6 signaling, apoptosis, inflammasome related regulators, and inflammation compared to IL-5 signaling, apoptosis, and basophil related processes associated with persistent carriers. This suggested that persistent carriers may have alterations in their ability to respond to Vita-PAMPs (pathogen associated molecular patterns) that may have facilitated persistent colonization with *S. aureus* (34, 49).

Vita-PAMPs are closely related to PAMPs in that they activate the innate immune system following recognition by the host’s pattern recognition receptors (PRRs), *e.g*. Toll-like receptors (TLRs), but differ in that they serve as microbial signatures of viability, *e.g*. bacterial mRNA, pyrophosphates, quorum sensing molecules, bacterial second messengers, and isopyrenoids resulting in heightened innate immune activation (34, 49, 50). For example, phagocytosis of live pathogens expressing both PAMPs and Vita-PAMPs, but not dead pathogens expressing PAMPs alone, resulted in the activation of a TRIF-dependent signaling pathway that triggered inflammasome activation and subsequent caspase-1 mediated production of INFβ (34). Activation of the inflammasome eventually results in a highly inflammatory form of cell death (pyroptosis) that is beneficial to pathogen containment/clearance since the release of endocytosed organisms results in more efficient clearance by neutrophils (51).

TLRs and other PRRs are the ‘guardians at the gate’ that define whether or not an immune response will be initiated, and depending on the nature of the stimulus these receptors will alter the type and quality of the adaptive response that develops. Ten TLRs have been described in humans and each TLR homodimer or heterodimer combination (*e.g*., TLR1:TLR2 and TLR2:TLR6) recognize unique PAMPs. All TLRs described to date utilize the MyD88/MAL signaling adaptor molecules with the exception of TLR4 that can also bind to the TRIM/TRAM adaptor molecule complex at the membrane of endosomal compartments (52); however, data suggested that TRIM/TRAM can also serve as an adaptor protein in TLR2:TLR6 signaling (53). Although TLR4 has long been recognized as the receptor for the Gram negative PAMP lipopolysaccharide (LPS) as well as a receptor for host-derived danger associated molecular patterns (DAMPs), it has also been shown to play a role in immune responses to Gram positive bacteria that generate lipoproteins and lipoteichoic acids typically recognized by TLR2 (52, 54). For example, TRAM-deficient macrophages infected with either herpes simplex virus or *S. aureus* displayed impaired type I interferon responses suggesting a role for TRAM in TLR2-dependent responses. Further complicating the TLR2 (Gram positive) TLR4 (Gram negative) dichotomy are data demonstrating that almost half of mice deficient in TLR4 died from an *S. aureus* infection suggesting that both TLR2 and TLR4 are important to abscess containment in addition to coordination of the T-cell response and mediating wound healing (55). In addition, TLR4 knockdown modulated the inflammatory response to *S. aureus* by down-regulating TNFα concentrations suggesting that both TLR2 and TLR4 contributed to the anti-*S. aureus* response (56, 57).

Our results suggested that although there are no or only few statistically significant differences between responses observed between persistent carriers and non-carriers following stimulation with either TLR ligands or bacteria, our data indicate that there may be potential alterations in the previously described Vita-PAMP pathway among *S. aureus* persistent carriers. These alterations appear to be localized to the TRIF/TRAM-related pathway suggested to be part of the recognition of live versus dead bacteria (58). The implications of these potential differences in TRIF/TRAM-related pathway activation between persistent carriers and non-carriers indicated that although the persistent carriers included in this study are fully capable of recognizing and responding to TLR ligands and dead bacteria, their responses to live bacteria are different and may be a determinant for persistent colonization with this organism.

## Materials and Methods

### Study population and sample collection

Pediatric patients at the Northwest Assistance Ministries Clinic, Houston, TX, located in northwestern Harris County were eligible for participation in the current study if they were participants of a prior parent study investigating pediatric *S. aureus* nasal carriage (n=438). In the parent study, we established nasal carriage phenotypes (persistent, intermittent, or non-carriers) based on the identification of *S. aureus* on 2 consecutive nasal swabs collected 12-17 days apart, based on the 2-culture method previously described (16, 17, 25, 59). Participants in the parent study testing positive for *S. aureus* on both swabs were classified as persistent carriers. Non-carriers were *S. aureus* negative at both time points and intermittent carriers were positive for one of the two initial swabs collected. No intermittent carriers are included in the present study. Using univariate statistics, we compared demographic characteristics between the two patient groups, using the chi-squared test for categorical variables (Fisher’s exact p-value if the comparison included a frequency ≤5) or ANOVA for continuous variables, using Stata 14 (Table 1) (College Station, TX).

Confirmatory third swabs were conducted on participants from the parent study until 15 persistent and 15 non-carriers of *S. aureus* were identified and also agreed to participate in the present study. Confirmatory third swabs were collected between 6-931 days after the second swab (median 207 days) (Table 2) and processed for the detection of *S. aureus* as described previously (16, 59, 60). *S. aureus* isolates from 12 of the 15 persistent carriers were available for analysis in the current study, and they were used individually (denoted “Carrier Strain”) and combined (denoted “Carrier Mix”). In all, a persistent carrier could have up to 6 *S. aureus* positive cultures from the 3 swabs collected at different time points (Table 2).

Peripheral blood was drawn into ACD Vacutainer^®^ tubes from participants enrolled in the study using standard phlebotomy techniques. Specimen tubes were labeled and transported at 4°C to the University of Texas Health Science Center at Houston School of Public Health for PBMC isolation.

### Ethics of experimentation

This study was approved by the Institutional Review Boards of the University of Texas Health Science Center and Benaroya Research Institute at Virginia Mason. Informed written consent was obtained from each participant’s parent or guardian. Child assent was obtained from all participants 7-11 years of age and adolescent consent from all participants ages 12-17 years.

### Targeted assay (TA) development

The TA was developed using a primarily data-driven approach. Thirty modules were selected based on how well they represented the signatures seen in the module analysis performed previously by our (33). The first principal component for each module was calculated and each probe was correlated to the first principal component for its respective module. Modules showing poor correlation between their first principal component and individual transcripts were removed from the selection process leaving 30 modules. The 3 best-correlated probes for the remaining modules were selected to represent the module in the TA (Table 3 and Table S1). The selected modules span a wide variety of cellular and immune functions including inflammation, the production of type I and II IFNs and other cytokines such as IL-1, IL-6, and IL-10, cytokines associated with apoptosis, caspase expression and FAS signaling, and cytokines associated with cell signaling/adhesion/chemotaxis. These functions are critical to the activation of innate immune responses. The mean, standard deviation, and coefficient of variation (cv) of the expression values for each probe across all donors were calculated and 6 probes with a low cv and means spanning the width of the expression values were selected as housekeeping genes (Table 3). In short, we used a data-driven approach to select 30 modules, 3 genes per module and 6 housekeeping genes that when combined, capture the same overall module signature as a full microarray or RNAseq (Table 3 and Table S1).

### Microarray analyses

Microarray analyses were carried out as part of the method development in order to compare the module signature between a full microarray and the TA. Frozen PBMCs from adult donors (n = 8, females, 21-60 years old) were treated as described in in the preceding section. The RNA from each sample was split and used for both microarray analysis and the TA as described above. Biotinylated cRNA targets were prepared from 250 ng of total RNA using the Illumina TotalPrep RNA Amplification Kit (Ambion, ThermoFisher Scientific). Labelled cRNA (750 ng) was then incubated for 16 h with HT-12 v4 BeadArrays (48,323 probes). Beadchip arrays were then washed, stained, and scanned on an Illumina HiScanSQ according to the manufacturer’s instructions.

Following background subtraction, raw signal values extracted with Illumina Beadstudio (version 2) were scaled using quantile normalization. Minimum intensity was set to 10 and all the intensity values were log_2_ transformed. Only the probes called ‘present’ in at least one sample (p<0.01) were retained for downstream analysis (n=19,624).

Transcripts differentially regulated upon stimulation were defined based on a minimum 2-fold change (up-or down-regulation) and a minimum absolute raw intensity difference of 100 compared to respective unstimulated samples. Heatmaps were generated using R (version 3.4.1). Networks were resolved with MetaCore™ (version 6.10, GeneGo).

### Validation of the TA

Correlations between the microarray and the TA platform were performed using the Spearman Rank Correlation on log_2_ FC values. Housekeeping values were compared using log_2_ intensity values and log_2_ delta CT values. To further validate the correlations, the correlation values were compared to 5,000 random permutations of all samples within each gene. The obtained permutated correlation values were compared to the correct correlation comparison and the number of permutated correlation values that was better than the real correlation value was added and then divided by the number of total permutations performed for each gene representing the p-value for the correlation of respective genes.

Correlations between individual modules and module clusters between microarray and TA were performed using the Spearman Rank Correlation on log_2_ FC values. Differentially-expressed genes and modules between PBMCs isolated from carriers and non-carriers stimulated with either live or dead bacteria were tested using the Student’s *t*-test and the Fligner-Killeen test was used to test variance. Differences between genes and modules between a commercial *S. aureus* strain, the unique carrier strain, and the mix of all unique carrier strains were tested using paired *t*-tests and the paired variance test. All p-values were False Discovery Rate (FDR) adjusted. All analyses were done in R (version 3.4.1).

### Test of Targeted assay robustness

The robustness of the TA was tested using whole blood, fresh PBMCs, and frozen PBMCs from the same blood draw from healthy adult volunteers (n = 4) at Benaroya Research Institute (Seattle, WA). From each blood sample (within 4 h of collection), 500 µl whole blood was diluted in 500 µl of RPMI containing respective stimuli in 48-well plates (Corning, Glenview, IL). The remaining blood was immediately used for PBMC isolation using Ficoll Paque plus (GE Healthcare, Littler Chalfont, U.K.). The isolated PBMCs were split in two, one half being used immediately and the other half frozen in 7% DMSO in fetal bovine serum and cooled at −1°C/min in a freezing container (Nalgene, ThermoFisher Scientific). Freshly isolated PBMCs were plated in 96 deep-well plates (Greiner Bio-One, Monroe, NC) at a concentration of 1×10^6^ cells/ml with a total of 0.5×10^6^ cells/well in RPMI 1640 (Thermo Fisher Scientific, Grand Island, NY) supplemented with 1X Serum Replacement (Sigma, St. Louis, MO),1X PenStrep (Sigma), 1 mM sodium pyruvate (Sigma), and the different stimuli used for PBMC activation.

Frozen PBMCs were thawed and plated in 96 deep-well plates (Greiner Bio-One, Monroe, NC) at a concentration of 1×10^6^ cells/ml with a total of 0.5×10^6^cells/well, in RPMI 1640 (Gibco-ThermoFisher Scientific) supplemented with 1X Serum Replacement (Sigma), 1X PenStrep (Sigma), and 1 mM sodium pyruvate (Sigma) and rested overnight at 37°C with 5% CO_2_ before adding respective stimuli.

PBMCs (freshly isolated and previously frozen) and fresh whole blood were each stimulated for either 2 or 6 h with (1) PMA/ionomycin (Sigma) (PBMC: 2.5 ng/ml PMA, 0.1 µg/ml ionomycin; whole blood: 25 ng/ml PMA; 1 µg/ml ionomycin, (2) IFNα (3×10^4^ units/ml, R&D Systems, Minneapolis, MN), (3) TNFα (20 ng/ml, R&D Systems), (4) IL-1β (PBMC: 2 ng/ml, whole blood: 20 ng/ml, R&D Systems), (5) PAM3CSK4 (10 ng/ml, InvivoGen, San Diego, CA), (6) poly (I:C) (25 µg/ml, InvivoGen), (7) LPS (100 ng/ml, InvivoGen), (8) R848 (300 ng/ml, InvivoGen), (9) CpGC (1.75 µM, InvivoGen), (10) heat-killed *Staphylococcus aureus* (10^7^ cells/ml, *Staphylococcus aureus* subsp. *aureus* Rosenbach (ATCC^®^ 6538™), or (11) un-stimulated for the same 2 or 6 h time period at 37°C with 5% CO_2_. Cells were then washed and lysed using RLT buffer (Qiagen, Germantown, MD) supplemented with 1% β-mercaptoethanol (Sigma) and whole blood was lysed with Tempus solution (Tempus Blood RNA tubes, ThermoFisher Scientific) to stabilize the RNA at 1:3 ratio. Cell/whole blood lysates were stored at −80°C until RNA extraction was conducted.

### Pediatric *S. aureus* carriage *in vitro* PBMC stimulation

Blood was collected at the Northwest Assistance Ministries Clinic (Houston, TX) and PBMCs were isolated from whole blood within 4 h of collection using Ficoll Paque plus (GE Healthcare) and frozen down in 7% DMSO in fetal bovine serum and cooled at −1°C/min in a freezing container (Nalgene, ThermoFisher Scientific). Frozen PBMCs were stored at −80°C before shipping on dry ice to Benaroya Research Institute for further processing and stimulation as described above.

Pediatric PBMCs were stimulated for 6 h with (1) TNF (20 ng/ml, R&D Systems), (2) heat-killed *S. aureus* (10^7^ cells/ml, InvivoGen), (3) *S. aureus* (10^7^ cells/ml) isolated from the subject’s own nasal swab (“Carrier Strain”) (*i.e*., only carriers were stimulated using their own carriage isolate), (4) a mixture (10^7^ cells/ml) of all *S. aureus* isolates obtained from carrier subjects (“Carrier Mix”), (5) IL-1β (2 ng/ml, R&D Systems), (6) PAM3CSK4 (10 ng/ml, InvivoGen), (7) CpG (1.75 µM, InvivoGen), or (8) un-stimulated, for the same time period. Due to the limited cell numbers that could be obtained from each pediatric donor, each treatment and control group/participant were carried out as singletons. After the incubation period, cells were washed with PBS and lysed using RLT buffer (Qiagen) supplemented with 1% β-mercaptoethanol (Sigma). Cell lysates were stored in −80°C until RNA extraction.

### RNA extraction and the targeted assay

RNA was extracted from PBMCs using the RNeasy Mini Kit (Qiagen) and from blood tempus mix tubes using MagMAX™-96 Total RNA Isolation Kit (Applied Biosystems, ThermoFisher Scientific). RNA integrity was assessed on an Agilent 2100 Bioanalyzer (Agilent, Palo Alto, CA).

After conversion of RNA to cDNA (50 ng total RNA from PBMCs and 100 ng total RNA from WB) (no globin reduction was performed) targeted genes were amplified and analyzed with Fast Gene Expression Analysis using EVAGreen^®^ on the BioMark™ HD System (Fluidigm, San Francisco, CA).

Raw CT values were exported from Fluidigm Real-Time PCR Analysis (version 3.1.3) and ΔCT and ΔΔCT values were calculated using R (version 3.4.1). Genes or samples that had a fail call rate of 50% or more were excluded from downstream analyses (all samples and genes passed this criteria in the experiments included in this study).

Transcripts differentially regulated upon stimulation were defined based on a minimum 2-fold change (up-or down-regulation) compared to respective unstimulated samples. Heat maps were generated using R (version 3.4.1).

### RNA sequence processing

RNAseq was performed on a representative subset of the samples from the respective *S. aureus* carriage phenotype groups. A total of 6 donors, 3 carriers (2 females and 1 male ages 4, 5, and 10 years, respectively) and 3 non-carriers (2 females, 1 male, ages 2, 4, and 11 years) treated with heat-killed *S. aureus* or a mixture of all collected carrier strains as described above (together with their untreated matched samples) were analyzed.

Sequencing libraries were constructed from total RNA using TruSeq RNA Sample Preparation Kits v2 (Illumina, San Diego, CA). Libraries were clustered onto a flowcell using a cBOT amplification system with a HiSeq SR v3 Cluster Kit (Illumina). Single-read sequencing was carried out on an Illumina HiScanSQ sequencer using a HiSeq SBS v3 Kit to generate 100-base single-end reads with a target of approximately 10 million reads per sample. The raw reads were processed in Galaxy, using TopHat (with bowtie) for alignment (GRCh37 as reference genome), BAM-to-SAM, Picard Alignment Summary Metrics, and Picard RNAseq Metrics. Genes were quantified using htseq-count. RPKMs for genes were obtained using edgeR (61, 62).

### RNAseq analysis

Raw counts were used for downstream analysis and Fragments Per Kilobase Million (FPKM) values were used for plotting. To identify differentially-expressed genes, a general linear model using negative binomial distribution with no random effects was applied using edgeR (61–63). Genes having an FDR <0.05 were considered differentially expressed.

Defining altered responses to live bacteria was done in one of two ways: (1) by applying the edgeR model with an FDR <0.05 as cutoff or (2) by calculating the dot (scalar) product and applying standard deviation (SD) cutoffs. All analyses and calculations were performed using R (version 3.4.1). Functional analysis and network enrichments of the differentially expressed genes and the genes identified as showing altered responses between live and dead bacteria were done using Pathway Studio ^®^ (version 12.0.1.9, Elsevier).

### Analysis of module-level data

The modules are designed based on the uniform gene expression of the genes within each module. Therefore, mmodule activity was calculated as the difference between the percent of up-regulated genes and that of down-regulated genes within each module. To obtain module results for the RNAseq data we first converted the Illumina microarray probe ID for each gene included in the modules to Ensembl gene IDs. Some probes shared Ensembl IDs and were dealt with as follows: i) multiple probes/Ensembl Gene IDs all in the same module were kept as one input and ii) multiple probes/Ensembl Gene ID in different modules can appear in multiple different modules. This was done to avoid picking the “best” fit. To determine if a gene was expressed or not in a particular module for the TA, fold change cutoffs were applied (log_2_ FC > ±2) and for the RNAseq data, the DEG cutoff FDR<0.05 was used.

## Data availability

The data described in this publication have been deposited in NCBI’s Gene Expression Omnibus (64) and are accessible through GEO Series accession number GSExxx (https://www.ncbi.nlm.nih.gov/geo/query/acc.cgi?acc=GSExxx).

## Supplementary Tables and Figures

**Supplementary Table 1.** Complete information about genes selected for the TA panel.

**Supplementary Table 2.** Spearman rank correlation values and permutation p-values for each gene present in both the targeted assay and the microarray.

**Supplementary Table 3.** Complete results from edgeR model for RNAseq analysis that is summarized in Table 5.

**Supplementary Table 4.** Complete dot product results and edgeR model results for comparison Carrier vs. Non-carrier for Live bacteria shown in Table 6 and Figure 9.

---

**Figure S1.**
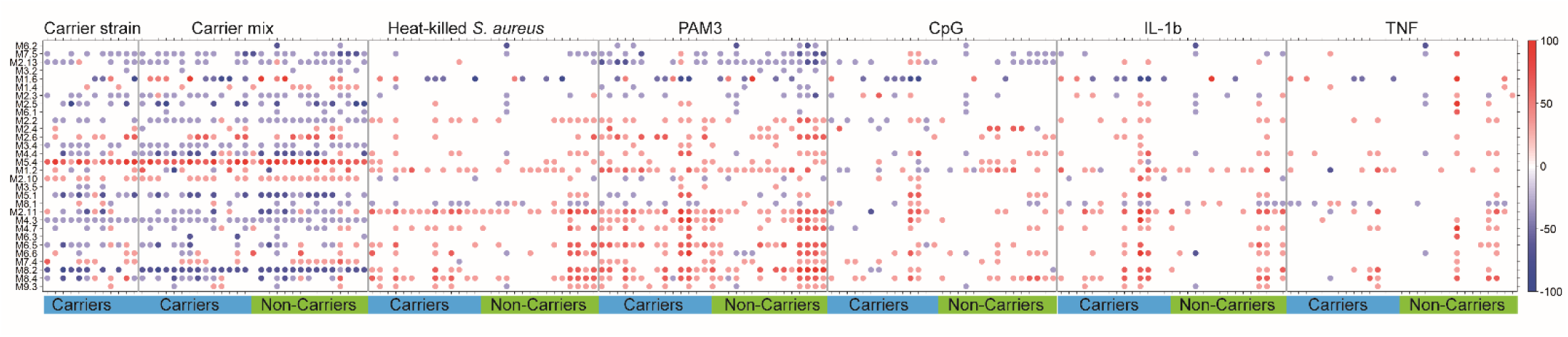
Module plots for pediatric PBMC stimulations

**Figure S2.**
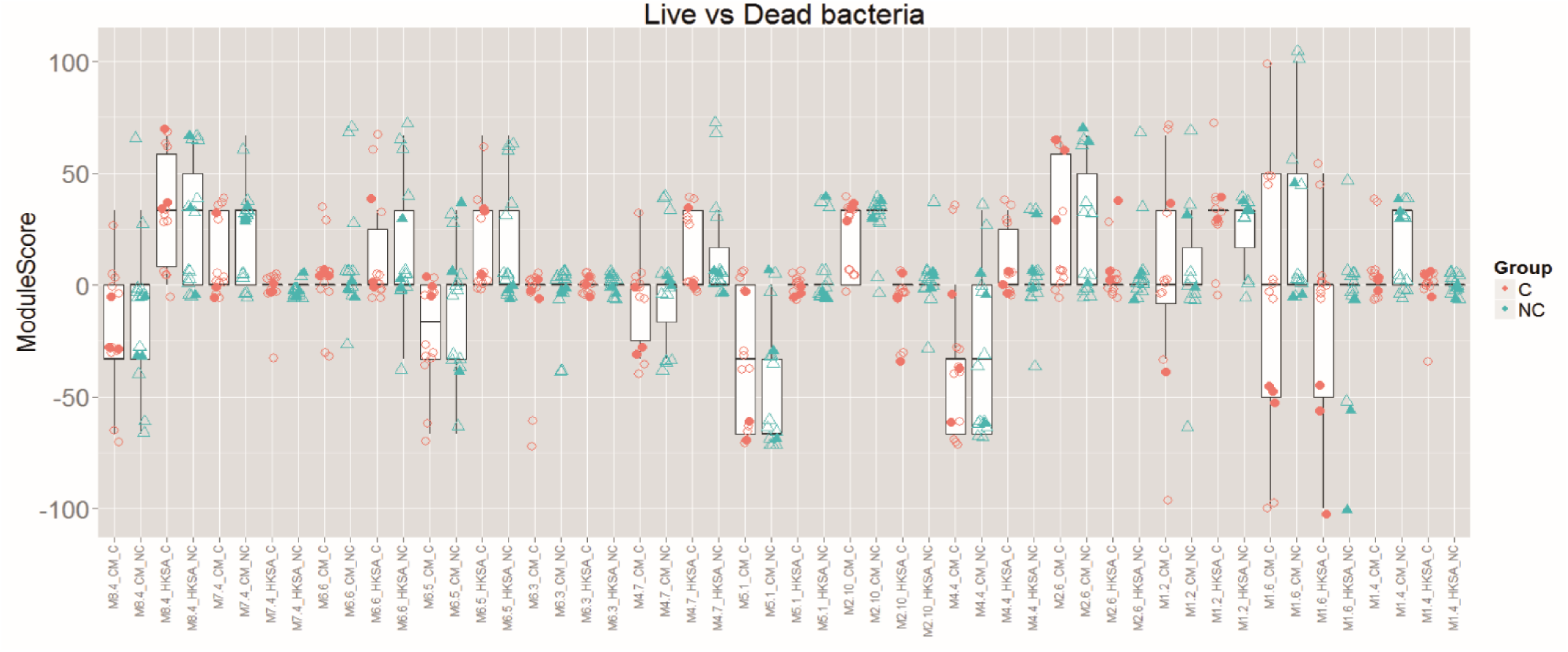
The donors selected for RNAseq. The selected donors represent the average response of the respective groups and are marked with filled symbols in the plot.

